# The phosphoproteomic landscape of the DNA damage response

**DOI:** 10.64898/2026.05.11.724232

**Authors:** Francesca Conte, Matthias Ostermaier, Juanjuan Wang, Christina Goss, Sergi Sayols, Jia-Xuan Chen, Vassilis Roukos, Frauke Gräter, Camilo Aponte-Santamaría, Katja Luck, Petra Beli

**Affiliations:** Institute of Molecular Biology (IMB), 55128 Mainz, Gemany; Department of General Biology, Medical School, University of Patras, Rio, Patras 26500, Greece; Max Planck Institute for Polymer Research, 55128 Mainz, Germany; Institute of Developmental Biology and Neurobiology (IDN), Johannes Gutenberg-Universität, 55128 Mainz, Germany

## Abstract

The DNA damage response (DDR) comprises an intricate network of protein–protein interactions and signaling pathways activated by DNA lesions and genomic instability. Central to this response is protein phosphorylation, which orchestrates DNA repair, cell cycle checkpoint activation, and chromatin organization. The response of the human phosphoproteome to different DNA damage-inducing agents and the functional role of regulated phosphorylation sites remains insufficiently characterized. Here, we systematically profiled the cellular phosphoproteome following exposure to eleven DNA damage–inducing agents that humans encounter physiologically or during cancer therapy. We identified a core set of DNA damage responsive phosphorylation sites, along with DDR signatures associated with DNA double strand breaks, replication stress and a pleiotropic response. Regulated phosphorylation sites are enriched within intrinsically disordered regions (IDRs), often forming clusters of nearby modifications that can affect IDR conformations or overlap with short linear interaction motifs. We discover that the RNA damage response predominantly shapes the changes induced by reactive aldehyde formaldehyde, alkylating agent methyl methanesulfonate and oxidative stress. Finally, we demonstrate that the proteasome-associated ubiquitin E3 ligase UBE3A is targeted by ATM and ATR kinases, thus linking proteasome regulation with the DDR.

## Introduction

Human cells are constantly exposed to DNA damage inducing toxins and chemicals from the environment or to the ones produced intrinsically as byproducts of metabolism. In addition, chemotherapeutic drugs and radiation therapy used for cancer treatment induce DNA damage. The DNA damage response (DDR) orchestrates a dynamic network of protein interactions and signaling events that safeguard genome integrity^1,2^. Protein phosphorylation mediated by kinases Ataxia telangiectasia mutated (ATM) and Ataxia telangiectasia and Rad3-related (ATR) plays a central role in the DDR^3^. ATM and ATR promote DNA repair by recruiting repair factors to damaged chromatin and trigger cell cycle checkpoints through activation of downstream checkpoint kinases CHEK1 and CHEK2^2,4^. Furthermore, the DDR regulates chromatin organization and nuclear architecture^4,5^. We have shown that RNA-binding proteins (RBPs) take part in the DDR by either being directly phosphorylated by ATM/ATR or by stress-induced kinases of the MAPK family including p38^6,7^. In addition to DNA damage, chemotherapeutic drugs and radiation therapy display pleiotropic effects on other cellular macromolecules such as RNA and proteins; however, the DNA damage-independent effects have so far been neglected. One example is the cellular response to ultraviolet light (UV-C) irradiation and formaldehyde (FA) and their toxicity, which has been attributed to the formation of transcription-blocking lesions in the genome. Recent studies, including our own work, reported the effects of UV-C and FA on RNA and proteins by induction of RNA-protein crosslinks (RPCs) that inhibit translation^8,9^, and demonstrated that UV-C-induced cell death is mediated by the ZAKα-dependent ribotoxic stress response (RSR)^10^. Similarly, 5-fluorouracil (5-FU), which is a common chemotherapeutic drug, induces RSR and cell killing in an RNA-dependent manner^11^.

To systematically define and compare how common DNA-damage inducers reshape the cellular phosphoproteome—and thereby resolve distinct DDR signatures—we conducted quantitative phosphoproteomic screens following eleven treatments, including replication stressors, double-strand break inducers, oxidative agents, and crosslinking compounds. We characterize phosphoproteome alterations elicited by diverse DNA-damaging agents and identify phosphorylation sites predominantly driven by DSBs, replication stress, or by DNA damage–independent, pleiotropic effects of these drugs on other cellular macromolecules. We demonstrate that regulated phosphorylation sites are enriched in intrinsically disordered regions (IDRs), which contain clusters of neighboring sites that modify charge distribution and induce changes in IDR conformation. We determine the predominant RNA damage-induced changes in the phosphoproteome response to FA, alkylating drug methyl methanesulfonate (MMS) and oxidative stress induced by H_2_O_2_. Furthermore, we demonstrate that autism- and proteasome-associated ubiquitin E3 ligase UBE3A is phosphorylated in an ATM/ATR-dependent manner, thus directly linking proteasome function with the DDR.

## Results

### Defining the regulatory phosphoproteome of the DDR

To elucidate the molecular signatures of the DDR induced by different DNA damage-inducing agents, we employed a tandem mass tag mass spectrometry (TMT-MS)-based approach for relative quantification of the phosphoproteome and total proteome in human osteosarcoma (U2OS) cells. U2OS cells were treated with chemotherapeutics (hydroxyurea [HU], gemcitabine [GEM], etoposide [ETO]), environmental toxins and radiations (methyl methanesulfonate [MMS], formaldehyde [FA], UV-C and X-ray), DNA replication inhibitor aphidicolin (APH) and oxidative stress-inducing agents (sodium arsenite [NaAsO_2_] and hydrogen peroxide [H_2_O_2_]) (Figure 1A, B). By mechanism of action, these can be grouped into topoisomerase poisons (ETO, CPT), replication stressors (APH, HU, GEM), oxidative stressors (H_2_O_2_, X-ray, AsO_2_), crosslinkers (FA, UV-C) and alkylators (MMS). To study the acute response, cells were treated with equitoxic doses of drugs for 2 hours and lysed immediately thereafter to avoid cell cycle-related effects. In case of X-ray and UV-C, cells were lysed 1- or 2-hours post-radiation, respectively (Figure 1A, S1A). Apart from AsO_2_ and FA, all treatments induced a significant increase in the phosphorylation of histone variant H2AX on S139 (γH2AX) that is targeted by ATM and/or ATR after DNA damage or replication stress, respectively (Figure S1B). Treatment of cells with ETO, CPT, H_2_O_2_, X-ray and MMS resulted in formation of DNA double strand breaks (DSBs) monitored by the chromatin-bound 53BP1 as proxy. Replication stressors (HU, GEM, APH), crosslinkers (UV-C, FA) and AsO_2_ did not induce relevant amounts of DSBs (Figure S1B). Even though cell cycle profiles remained unchanged for all treatments (Figure S1C), DNA replication was inhibited to various degrees, apart from X-ray irradiation that did not influence the amount of incorporated thymidine analogue 5-Ethynyl-2′-deoxyuridine (EdU) (Figure S1C).

**Figure 1.**
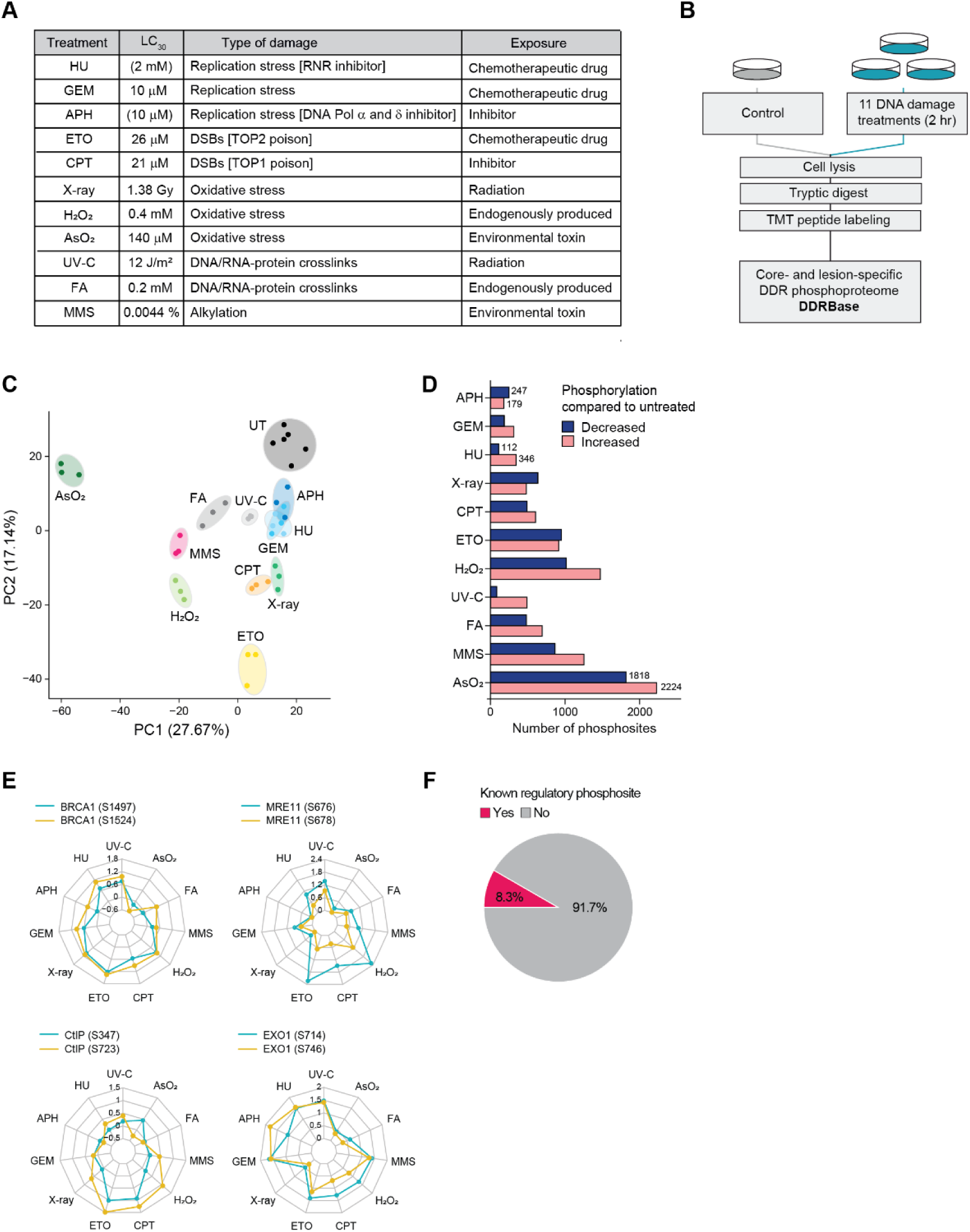
Defining the regulatory phosphoproteome of the DDR. A. Summary of agents used in this study, their mechanism of action, and treatment conditions for the phosphoproteome analysis. B. Experimental scheme of TMT-based quantitative phosphoproteomics after treatment of human osteosarcoma (U2OS) cells with DMSO or eleven DNA damage-inducing treatments (n=3) Cells were treated for 2 hours with the selected DNA damaging treatments and immediately harvested for cell lysis. For irradiation treatments, cells were exposed to the respective agent and allowed to recover for 2 hours (X-ray) or 1 hour (UV-C) prior to harvest and lysis. C. Principal component analysis (PCA) of analyzed DDR phosphoproteomes. D. Bar graph showing the number of significantly regulated phosphorylation sites for each treatment (FDR ≤ 0.05; absolute fold-change > 1.5). All regulated sites can be found in Table S1. E. Radar plots showing the log_2_(fold-change) of selected previously characterized DDR-induced phosphorylation sites. F. Pie chart showing the percentage of significantly regulated phosphorylation with known function (obtained from the PhosphoSitePlus PTM database).

Principal component analysis (PCA) was used to identify similarities and differences in the phosphoproteome between responses: AsO_2_ clustered away from other treatments, suggesting a more diverse effect on the phosphoproteome (Figure 1C). This was also true but to a lesser extent for other pleiotropic agents that induce protein and RNA damage in addition to DNA damage (H_2_O_2_, MMS and FA) (Figure 1C). We identified 2,467 proteins harboring 6,070 phosphorylation sites that changed significantly in at least one treatment (Table S1). Pleiotropic treatments including AsO_2_, H_2_O_2_ and MMS showed the most pronounced effect on the phosphoproteome, leading to significantly increased phosphorylation of 15%, 10% and 8% of all quantified phosphorylation sites, respectively. On the other hand, replication stressors APH, HU and GEM showed a milder response, with increased phosphorylation of less than 3% of quantified phosphorylation sites (Figure 1D). Notably, we recapitulated phosphorylation events with known functions in the DDR: for instance, homologous recombination (HR) factors showed increased phosphorylation on known regulatory sites upon DSB-inducing agents, including S1497 and S1524 of BRCA1 (regulating BRCA1 foci formation at DSBs)^12–16^, S678 of MRE11 (modulating the binding of MRE11 to chromatin)^17–19^ and S347 of RBBP8/CtIP (regulating CtIP endonuclease activity)^20^ (Figure 1E). Treatment-specific regulation was also recapitulated: S746 phosphorylation on DNA repair factor EXO1, which is required for the protection of stressed replication forks from unscheduled fork resection^21,22^, displayed the strongest increase in response to replication stressors, as opposed to S714 that more generally regulates EXO1 stability^23^ (Figure 1E). Notably, functional characterization was present for only 8.3% of the significantly regulated phosphorylation sites (as annotated in the PhosphoSitePlus PTM Database)^24^, highlighting the need for systematic approaches to uncover the regulatory mechanisms of the DDR (Figure 1F). Among regulated sites, 1,083 displayed a functional score^25^ > 0.5, suggesting that these sites likely have a molecular function based on a collection of 59 parameters including residue conservation, structural properties and regulation by upstream kinases (Table S1).

### Protein kinase-substrate relationships in the DDR

We performed kinase activity prediction based on Kinase-Substrate Enrichment Analysis (KSEA)^26^ to determine kinases targeting DNA damage-responsive phosphorylation sites (Figure 2A; Table S1). The PIKK kinases ATM, ATR, and PRKDC (DNA-PK) were activated by all treatments except for AsO_2_; this was substantiated by motif analysis of induced phosphorylation sites, which showed the expected PIKK-specific S/T-Q motif (Figure 2B, C). On the contrary, activity of cell cycle-regulating cyclin-dependent kinases CDK1 and CDK2 was inhibited by all treatments (Figure 2A). While replication stress- and DSB-inducing treatments primarily led to the activation of PIKK kinases and direct downstream kinases CHEK1 and CHEK2, H_2_O_2_, MMS, FA and AsO_2_ activated a wider range of kinases, indicative of their impact on additional cellular processes. This activation pattern was shared by mitogen-activated protein kinases (MAPKs), as well as members of the RPS6KA/B, AKT, and PKC families (Figure 2A). In line with the DNA damage-dependent activation of cell cycle checkpoints, most decreased phosphorylation sites were found within the S/T-P-X-R/K motif, which is targeted by CDK1/2 (Figure 2A, S2A). The largest absolute number of induced S/T-Q phosphorylation sites were found after DSB-inducing treatments ETO, CPT and H_2_O_2_ (Figure 2C). On the contrary, AsO_2_ led to the strongest perturbation of the phosphoproteome, but displayed the smallest fraction of induced S/T-Q phosphorylation sites (only 2% of induced sites conformed to the S/T-Q motif compared to 38% in case of UV-C and GEM that induce replication stress) (Figure 2C). Gene Ontology (GO) enrichment analysis of proteins harboring increased S/T-Q or S/T-P phosphorylation sites revealed two different sets of proteins: S/T-Q phosphorylation sites were found predominantly on DNA repair factors and nuclear inclusion body proteins, while phosphorylation of S/T-P sites (where we excluded sites that were predicted to be targeted by CDK1/2) occurred on proteins involved in MAPK signaling, GTPase regulation and cytoskeleton protein-binding, linking those sites to general MAPK-dependent stress response (Figure 2D). The degree of conservation of a phosphorylation site across species has been shown to be a good proxy for biological function^27–29^. To address phosphorylation site conservation, we obtained the relative local conservation (RLC) score in metazoa for all quantified sites using PepTools^30,31^ (Table S2). We then sorted DNA damage-responsive sites based on their presence within an S/T-Q, S/T-P, or S/T-P-X-R/K motif (sites matching to both S/T-P and S/T-P-X-R/K were assigned to the latter category), and evaluated their conservation (Figure S2B). We found S/T-P-X-R/K phosphorylation sites (CDK1/2 targets) to be the most conserved across categories (Figure S2B). Notably, regulated S/T-Q phosphorylation sites were less conserved than S/T-P sites; this is in line with previous reports showing the presence of the S/T-P motif across diverging eukaryotic lineages, which indicates an early origin in eukaryotic evolution, as opposed to the predominantly metazoan S/T-Q motif ^32^. Furthermore, regulated S/T-Q phosphorylation sites exhibited lower overall conservation than non-regulated sites; notably, this difference was no longer apparent when conservation was assessed specifically within mammals (Figure S2C).

**Figure 2.**
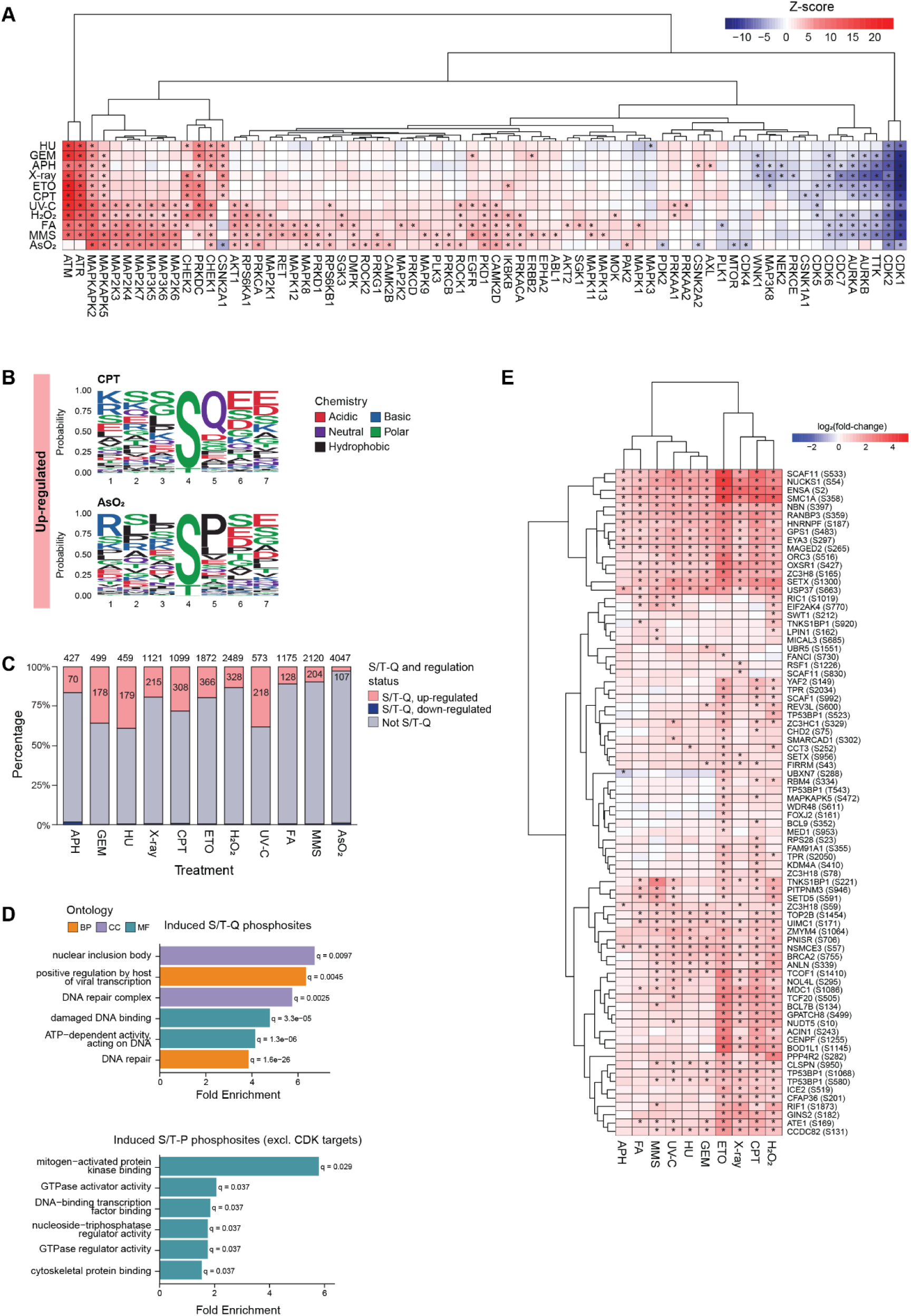
Protein kinase-substrate relationships in the DDR. A. Heatmap displaying Kinase-Substrate Enrichment Analysis (KSEA) for the DDR phosphoproteome. B. Sequence motif analysis of CPT or AsO_2_-specific phosphorylation sites with IceLogo, matching the S/T-Q and S/T-P motifs, respectively. Sequence windows of significantly upregulated phosphorylation sites were compared to all quantified phosphorylation sites. Amino acid chemistry is indicated (right). C. Bar graph showing the percentage of S/T-Q sites with significant increase (FDR ≤ 0.05; fold-change ≥ 1.5), decrease (FDR ≤ 0.05; fold-change ≤ -1.5), or no change in abundance for each treatment. D. Gene Ontology enrichment analysis of proteins with increased phosphorylation on the S/T-Q motif (upper panel), or of increased phosphorylation on the S/T-P motif where proteins with CDK1/2 motif were excluded (lower panel). AsO_2_-regulated sites were excluded from this analysis. E. Heatmap of non-functionally characterized and highly conserved S/T-Q sites that showed significantly increased phosphorylation in at least one treatment (FDR ≤ 0.05; fold-change ≥ 1.5).

Phosphorylation of S/T-Q sites is likely executed by ATM/ATR, linking these regulatory events directly with the DDR. Thus, we categorized induced S/T-Q sites based on their conservation into sites with high, medium and low conservation (Figure S2D). Highly conserved sites (low RLC score; 99 sites) were present on *bona fide* DNA repair proteins such as CHEK1, BRCA1, MRN complex, ATR and ATM (Table S3). Surprisingly, only 12 out of the 99 highly conserved sites are annotated as functionally characterized, highlighting the substantial opportunity to uncover new regulatory mechanisms within the DDR (Figure 2E). Overall, 45 S/T-Q sites identified in our screen have known functions; however, only 22 of these exhibit an RLC score < 0.5 (Table S3). For instance, less conserved and previously functionally characterized S/T-Q sites are present on FANCA/FANCD2^33,34^, EXO1^23^, VCP^35–37^, CHD4^38^ and PARP1^39^ among others (Table S3). Interestingly, lowly conserved S/T-Q phosphorylation sites are present on “damaged DNA binding” proteins, but also proteins annotated with GO terms not directly related to DNA repair such as “protein-DNA complex” and “U2-type spliceosomal complex” (Figure S2E; Table S3). These include sites on transcription and RNA processing regulators including p300, RUVBL2, BRD8, VPSS72, HTATSF1, SF3A1, SNW1, SF3B2 and SNIP1 (Figure S2E, F; Table S3). Taken together, our findings demonstrate that the functional scope of ATM/ATR signaling has broadened to regulate additional DNA repair proteins and chromatin-associated processes such as transcription and pre-mRNA processing in response to DNA damage, and that sequence conservation of S/T-Q phosphorylation sites is not a reliable indicator of their functional relevance in human cells.

### Specific and common responses of the phosphoproteome to DNA damage-inducing agents

To identify specific signatures of the DDR-induced phosphoproteome, we combined self-organizing map (SOM) clustering with k-means analysis of all significantly regulated phosphorylation sites (Figure 3A, Table S4). AsO₂-induced sites were excluded because this treatment caused widespread phosphoproteomic remodeling. The DSB-inducing agents (X-ray, ETO, CPT) clustered together, as did replication stressors (HU, APH, GEM), revealing distinct phosphorylation signatures for these two types of genotoxic stress (Figure 3A). Pleiotropic treatments (FA, MMS, H₂O₂) also clustered together, pointing to a shared stress response pathway that operates independently of DSB and replication stress responses. UV-C did not cluster with other treatments but showed the strongest resemblance to replication stressors (Figure 3A).

**Figure 3.**
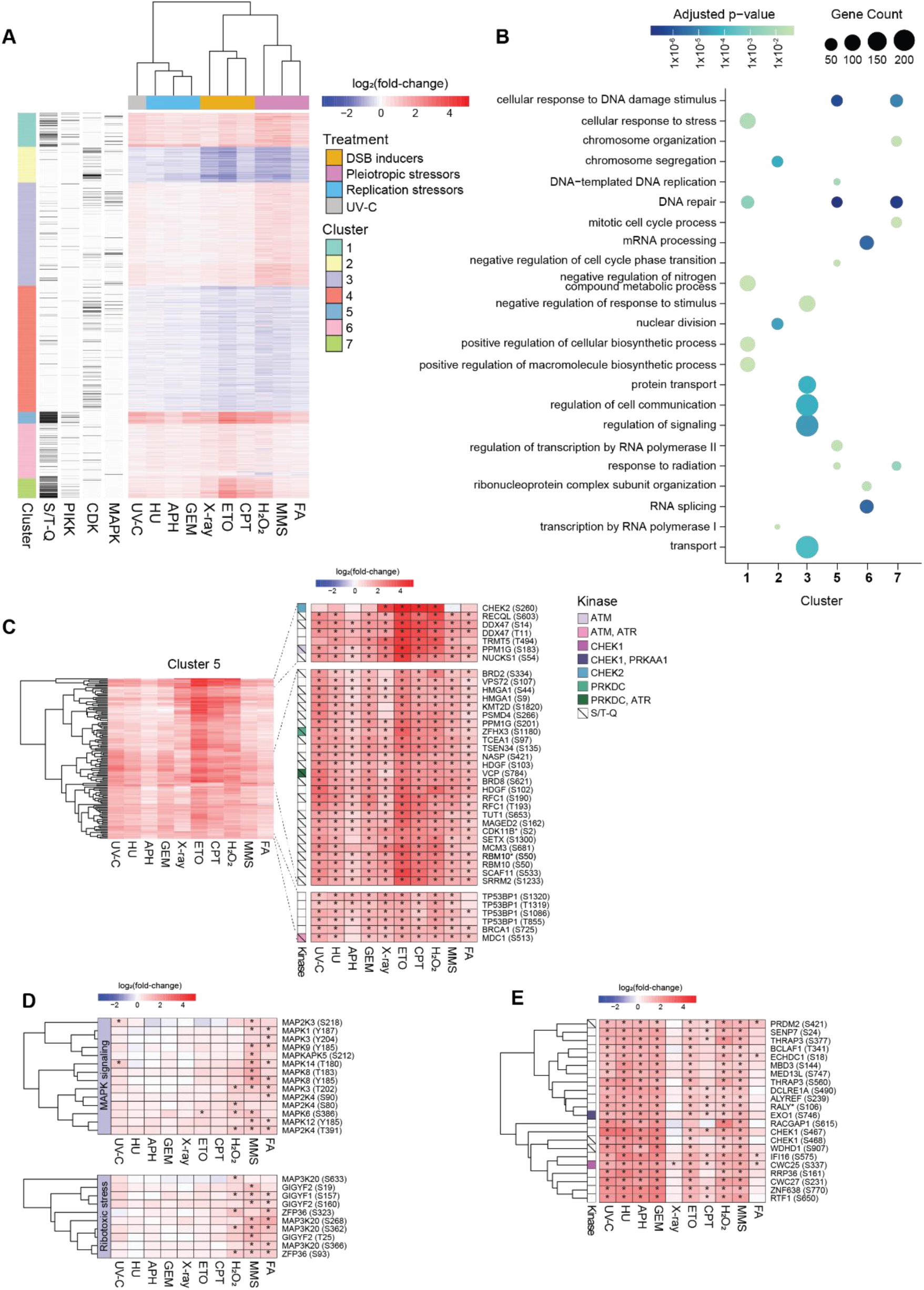
Specific, common, and pleiotropic responses of the phosphoproteome to DNA damage-inducing agents. A. Heatmap of phosphorylation sites significantly changing (FDR ≤ 0.05; absolute fold-change > 1.5) in at least one condition (excluding AsO_2_). Clusters obtained using Self-Organizing Maps (SOM) followed by k-means (k=7) are shown. Phosphorylation sites are cross-referenced with the results of the KSEA analysis (CDKs, MAPKs, and PIKKs substrates) and S/T-Q sites are annotated. B. Dot plot showing the comparative Gene Ontology term (biological process) enrichment analysis for clusters identified in figure 3A. Only clusters with significant gene ontology terms (BH-adjusted p-value < 0.05) are shown. When possible, the first five terms by fold enrichment per cluster are shown. Dot size corresponds to gene count. C. Heatmap showing all cluster 5 phosphorylation sites (left), or zoomed-in panels with sites of interest (right). Phosphosites are cross-referenced with the results of our KSEA analysis (CDKs, MAPKs, and PIKKs substrates) and S/T-Q sites are annotated. D. Heatmaps recapitulating phosphorylation events in the MAPK pathway (upper panel) and the ribotoxic stress response (lower panel). E. Heatmap showing a selection of phosphorylation sites from Cluster 1. Phosphorylation sites are cross-referenced with the results of our KSEA analysis (CDKs, MAPKs, and PIKKs substrates) and S/T-Q sites are annotated.

Based on the phosphorylation patterns, we defined clusters that are primarily driven by DSB signaling (cluster 6, 7), common DSB and replication stress signaling (cluster 5) and by a pleiotropic response (cluster 1, 3). Cluster 5 displayed the strongest and most homogenous response, with 48 out of 129 phosphorylation sites undergoing a significant increase in their levels across all treatments (Figure 3A). Most phosphorylation sites in cluster 5 were found within an S/T-Q motif (79%) (Figure 3A) and on proteins annotated with GO terms “DNA repair”, “response to radiation” and “regulation of transcription by RNA polymerase II” (Figure 3B). This cluster included subunits of protein complexes involved in DNA repair (MRN complex - NBN1, MRE11), chromatin remodeling (Nuclear mitotic cohesin complex - SMC1A, SMC3) and transcription (SNIP1/SkIP associated RNA-processing complex - SNW1, SNIP1, PNN, BCLAF1, THRAP3) (Figure S3A). Proteins previously not linked to or only loosely associated with the DDR such as KMT2D, DDX47, RBM10 and ZFHX3 were also found in this cluster (Figure 3C; Table S4), highlighting the potential of comparative phosphoproteomics screens for discovering new genome maintenance factors. In summary, proteins belonging to cluster 5 contain phosphorylation events that are part of a core response to DSBs and replication stress primarily targeted by the DDR kinases ATM/ATR (Figure 3C; Table S4).

Clusters 3 and 7 showed opposite trends, including phosphorylation sites that responded more strongly to pleiotropic agents or DSB-inducing agents (and H_2_O_2_), respectively (Figure 3A, S3B). Cluster 7 (DSB response) showed enrichment for DNA repair and chromosome organization terms, driven by the increased phosphorylation of subunits of chromatin-associated complexes (SWI/SNF ATP-dependent chromatin remodeling complex - SMARCA2, SMARCC1, SMARCE1; GINS complex - GINS2, GINS3; general transcription factor TFIIIC complex - GTF3C1, GTF3C3, GTF3C4) (Figure S3A, B, Table S4). Proteins involved in the regulation of chromosome organization displayed increased phosphorylation after DSBs but not after replication stress (Figure 3B, S3B). These included sites on subunits of the BAF chromatin remodeling complex such as SMARCC1, PBRM1, SMARCE1, SMARCAD1 and SMARCA2 (Figure S3A, B).

Conversely, differentially phosphorylated proteins in cluster 3 (pleiotropic response) are associated with processes that are non-exclusively nuclear, such as “regulation of signaling”, “transport” and “regulation of cell communication” (Figure 3B). This response was driven primarily by MAPKs and AKTs kinases and included numerous phosphorylation events on the MAP/ERK kinase pathway (among these: MAPK3/ERK1 - T202 and Y204; MAPK1/ERK2 - Y187; MAPK8/ JNK1 -T183 and Y185; MAPK14/p38 - T180; MAP3K20/ZAKa - S268, S362, S366, S633) and ribotoxic stress factors (Figure 3D). We conclude that the pleiotropic response of H_2_O_2_, FA and MMS (encapsulated in cluster 3, Table S4) is primarily driven by MAPK signaling because of the ribotoxic stress response and is independent of DNA damage-inducing effects of those treatments. Interestingly, we found that 13.6% of phosphorylation sites in cluster 3 were predicted targets of CDK1 and, contrary to the inhibition of CDKs observed across treatments, were induced in response to H_2_O_2_, MMS or FA (Figure S3C). Motif analysis showed that these sites fulfilled the minimal S/T-P CDK1 motif, which differed from the canonical motif S/T-P-X-R/K present in CDK1/2 targets in cluster 2 and cluster 4 (Figure S3C, D). These included previously described CDK1 targets, such as serine 65 and threonine 59 on UBXN2B^40^, serine 221 on NUP50^41,42^ and serine 259 on PIK3C2A^43^ (Figure S4C).

Replication stress-specific signaling was scarce and primarily overlapped with DSB response; however, cluster 1 contained a subset of proteins that were regulated more strongly after replication stress compared to DSBs (Figure 3E). Apart from the ATR downstream kinase CHEK1 and the replisome component WDHD1, this subset contained RNA processing factors including ZNF638, CWC27, RALY, ALYREF, THRAP3 and BCLAF1 (Figure 3E). These proteins did not show increased phosphorylation after X-ray exposure, in line with the observation that their regulation is predominantly a consequence of perturbed S-phase and replication stress (Figure 3E).

Lastly, clusters 2 and 4 included the majority of dephosphorylation events, driven by the inhibition of CDKs (respectively 65% and 45% of sites were predicted as CDK substrates) (Figure 3A, S3C). Strikingly, nine phosphorylation sites in cluster 2 were uniformly decreased across all examined treatments: these included histone H1-4 (T18), transcriptional regulators E2F7 and MTA1 (S95; T564 and S576), replication factors MCM10 and ORC1 (S155; S287), cohesin accessory factor PDS5A (S1206 and T1208), and nuclear pore complex subunit NUP153 (T699). The differentially phosphorylated proteins in these clusters function in cell cycle regulation and associated processes (Figure 3B); for instance, we observed loss of phosphorylation of subunits of the kinetochore CCAN complex (CENPC, CENPN, CENPQ, CENPT), nuclear origin recognition complex (ORC1, ORC2) and B-WICH chromatin remodeling complex (BAZ1B, SMARCA5, SF3B1, MYBBP1A, DEK, DDX21) (Figure S3A).

### DDR-induced multi-site cluster phosphorylation regulates conformational dynamics of IDPs

To further characterize the DNA damage-responsive phosphorylation events, we investigated the biophysical properties of phosphorylation sites. We found that the average accessibility based on AlphaFold (namely, the proportion of the phosphorylated peptide that is accessible to water molecules) and IUPred scores for regulated sites were respectively 0.775 and 0.69, demonstrating that DNA damage-responsive phosphorylation sites are enriched in regions of high solvent accessibility and intrinsic disorder. This aligns with previous studies showing that phosphorylation predominantly occurs in intrinsically disordered regions (IDRs)^44,45^. Only 2.3% of all quantified phosphorylation sites were localized in structured regions, which were, interestingly, enriched in catalytic and calcium-binding domains (Figure S4A, B; Table S5).

Phosphorylation can appear in clusters of sites, with multiple nearby residues being phosphorylated cooperatively by the same or distinct kinases^46,47^. We identified a total of 1,205 phospho-clusters on 540 proteins, involving 36.6% of the DNA damage-responsive phosphorylation sites (Figure 4A; Table S6); the number of phospho-clusters correlated with the number of regulated sites per treatment, with pleiotropic treatments (with the exception of FA) registering the highest number of phospho-clusters, followed by DSB inducers and replication stressors (Figure S4C). Specifically, we found 34 and 36 phospho-clusters to be fully gained or lost in at least one treatment (namely, phospho-clusters composed only of up-regulated and down-regulated phosphosites), respectively (Figure 4B, S4D). Overall, the phosphorylation pattern of phospho-clusters (the mixture of regulated and non-regulated sites) was treatment-specific, with limited overlap between treatments with the same mode of action (e.g. only 12 and 10 phospho-clusters shared the same regulation between DSBs agents ETO and CPT, and ETO and X-ray, respectively) (Figure S4E). Only one phospho-cluster displayed the same regulation across all treatments: this included three phosphorylation sites on DDX47, a poorly characterized RNA helicase with roles in pre-mRNA processing.

**Figure 4.**
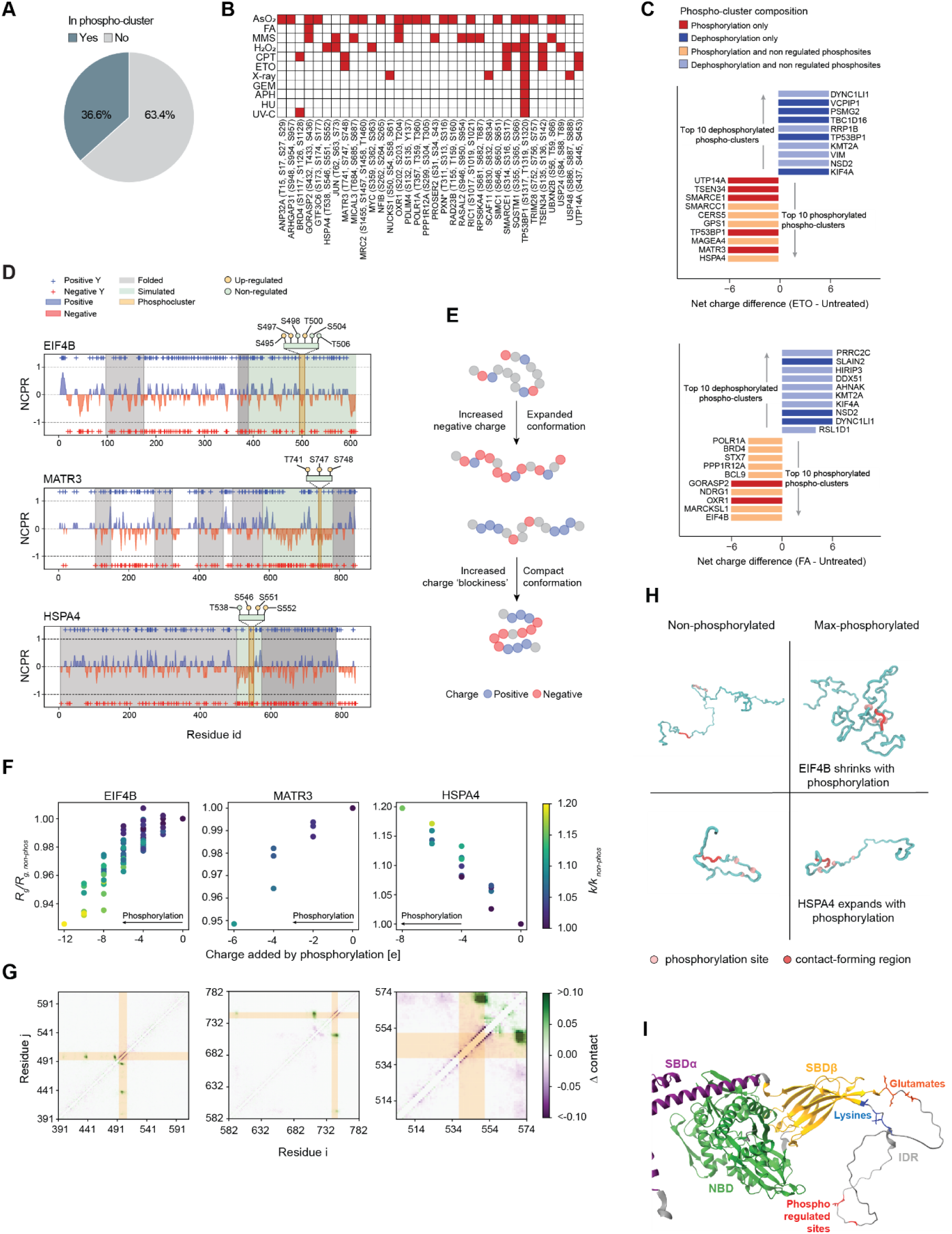
DDR-induced multi-site cluster phosphorylation regulates conformational dynamics of IDPs. A. Venn diagram showing the percentage of DNA damage-responsive phosphorylation sites (FDR ≤ 0.05; absolute fold-change > 1.5) within phospho-clusters. Phospho-clusters were defined as groups of two or more phosphorylated residues within a 10-amino acid stretch that contained at least two DNA damage-responsive sites. B. Heatmap showing the enrichment of phospho-clusters fully gained upon DNA damage treatments. An asterisk next to a gene name indicates an isoform other than the canonical one. C. Ranked bar plots displaying the difference in net charge (total charge treatment - total charge untreated) for individual phospho-clusters upon selected treatments. The top 10 phospho-clusters acquiring a positive or negative charge are represented D. Charge plots showing the net charge per residue (NCPR) calculated along the protein sequence of EIF4B (upper panel), MATR3 (middle panel) and HSP4A (lower panel) (blue: positive and red: negative). Folded domains (predicted with AlphaFold), disordered regions, and phospho-clusters of interest are shown in grey, green, and orange, respectively. The + symbols indicate the exact position of charged amino acids. E. Cartoon showing the effects of an increase in negative charge (left) or in charge “blockiness” (right) on the conformation of a stretch of amino acids. Neutral amino acids are depicted in grey. F. Scatter plot showing the average radius of gyration, normalized by that of the corresponding unphosphorylated fragment, (R_g_/R_g,non-phos_) plotted as a function of the negative charge added through phosphorylation, for EIF4B (left), MATR3 (center) and HSPA4 (right) fragments. The color indicates the patterning parameter κ, normalized by that of the corresponding unphosphorylated fragment (κ/κ_non-phos)_. See the absolute values of the R_g_ in Figure S5A. G. Contact maps illustrate the contact probability (Δ contact) for a given residue pair (i,j), for EIF4B (left), MATR3 (center) and HSPA4 (right) fragments. A Δcontact >0 (in green) indicates a more frequent contact, in the fully phosphorylated fragment versus the unphosphorylated version, and Δcontact <0 (in purple), indicates the opposite behavior. Phospho-cluster location is indicated in orange. H. Example conformations of two protein fragments illustrate how phosphorylation-mediated contact formation can lead to different size trends, i.e. either compaction (top row: shown here for EIF4B, but equivalent for MATR3) or extension (bottom row: HSP4A). The phosphorylation sites are highlighted as pink spheres and the regions engaging in contacts with the phospho-cluster as red ribbons. Representations are not at scale. I. AlphaFold-generated structural model of human HSPA4^54^. The IDR used in simulations is shown in grey. Phosphoregulated sites are indicated in red. The positively charged residues at the C-terminus of the IDR (labelled as lysines) that establish contacts in simulations with the phosphoregulated residues are indicated. In unphosphorylated state, these lysines were observed in simulations to establish contacts with the highlighted glutamates. The IDR is inserted between the second last and last β-strand of the SBDβ domain. This massive loop appeared to arise during vertebrate evolution. Domain coloring according to TED predictions^55^

As the addition of a phosphate group confers a double negative charge at physiological pH (pKa ∼ 6.7), the local accumulation or loss of phosphorylation sites can respectively increase or decrease the fraction of charged residues and net charge per residue (NCPR) of a phospho-cluster. Indeed, numerous phospho-clusters underwent remarkable changes in absolute charge, underlying potential changes in the protein’s local biophysical properties (Figure 4C; Table S7). To investigate how phosphorylation impacts conformational dynamics, we selected three poorly characterized proteins in DDR - EIF4B, MATR3 and HSPA4 – for in-depth simulation analysis. These proteins harbor phospho-clusters exhibiting a large change in net charge upon FA (EIF4B) and etoposide (HSPA4, MATR3) treatment (Figure 4C). As expected, the phospho-clusters were found in IDRs of the respective proteins, interestingly, featuring already in unchallenged cells a pronounced level of negative charge clustering (Figure 4D, S4F). An increase in negative charge, such as the one caused by DNA damage-induced phosphorylation, was previously reported to correlate with the expansion of IDRs^48,49^. On the other hand, increases in phosphorylation can also lead to a segregation of positive and negative charges (also known as “charge blockiness”, *κ*) resulting in a more compact conformation of the IDRs^50^ (Figure 4E). To test which of these concepts better explain the conformational behavior of the three IDRs upon phosphorylation, we simulated the dynamics of these IDRs with stochastic molecular dynamics simulations for increasing levels of phosphorylation. We then correlated the change in radius of gyration of each IDR with increases in negative charge and charge blockiness expressed as *κ*. Intriguingly, we observed that the IDRs of EIF4B and MATR3 became more compact as negative charge was added through phosphorylation in line with a strong increase in *κ* (Figure 4F, G). In sharp contrast, increasing phosphorylation caused HSP4A to adopt a more extended conformation (higher R_g_), aligning with expectations based on added negative charges to the IDR but deviating from predictions based on charge blockiness (Figure 4F, G). Note that δ_-+_, a sequence-length independent charge-clustering parameter^51,52^ did not explain the observed trend either (Figure S4G). We have previously observed that the formation of specific intramolecular contacts upon phosphorylation mediates changes in the radius of gyration of IDRs^49^. Indeed, in all three simulated IDRs, we found that full phosphorylation led to the formation of new intramolecular contacts between residues of the phospho-clusters and other residues of the IDRs (Figure 4G). These contacts explain the compaction of the EIF4B and MATR3 IDRs but, interestingly, involve an extension of the HSP4A IDR (Figure 4H). According to our simulations, upon phosphorylation of HSP4A, its positively charged C-terminal end switches from interacting with a very N-terminal region of the IDR to interacting with the close-by phospho-cluster, leaving the N-terminal part of the IDR free to explore more extended conformations (Figure 4G, H). HSP4A is a member of the Hsp70 protein family. Structural studies done for other family members described intramolecular interactions between the folded nucleotide-binding domain (NBD) that is upstream of the simulated IDR and the downstream folded substrate-binding domain (SBD) corresponding to the open conformation of Hsp70 proteins^53^. Such intramolecular contacts between both domains are also predicted by AlphaFold for the full-length monomer model of HSP4A (Figure 4I). Based on our results, we propose that clustered phosphorylation of the linker IDR modulates the intramolecular contacts formed between the NBD and SBD domain, thereby regulating substrate-binding of HSP4A. Overall, results from all three simulated IDRs demonstrate that phosphorylation-induced conformational changes cannot be uniquely attributed to simple biophysical parameters such as net charge and charge distribution, but also to the formation and depletion of specific intramolecular contacts.

### DNA damage-induced phosphorylation sites regulate SLiM-based interactions

The activation of lesion-specific DNA repair pathways relies on networks of protein-protein interactions (PPIs) that regulate protein localization, activity and stability. Transient and low-affinity interactions are often mediated by short linear motifs (SLiMs), stretches of < 10 amino acids typically found in IDRs ^56,57^. Notably, phosphorylation of SLiMs plays a key regulatory role, modulating affinities and allowing for the creation of binding sites for obligate phospho-binding proteins^58,59^. To gain insight into the impact of DNA damage-dependent phosphorylation on PPIs, we investigated the overlap between SLiMs and differentially regulated phosphorylation sites. A list of experimentally validated SLiMs was obtained from PepTools, encompassing entries from the Eukaryotic Linear Motif (ELM) database and from the Motif Map of the Proteome (MoMaP)^60^. We identified 161 SLiMs harboring DNA damage-responsive phosphorylation sites, which were assigned to 7 motif types (“Cleavage”, “Degradation”, “Docking”, “Scaffolding”, “Modification”, “Trafficking”, “Unassigned”) (Figure 5A; Table S2). Of these, 61 phosphorylation sites were found to overlap with scaffolding motifs, which are involved in noncatalytic interactions with their respective binding domains (Figure 5B). As expected, several phosphorylation sites were found in motifs that function in DNA repair, such as the ones recognized by proteins containing BRCT domains (BRCT_phosphopocket and BRCT_TOPBP1; H2AX - S140/143, TP53BP1 - S366), FHA domains (NIFK - T238), and by the replication-associated protein PCNA (PCNA_PIP box; FAM111A - S26)^61–64^. The most represented scaffolding motif class was the 14-3-3 domain-binding consensus motif (22 out of 61 SLiMs), which requires phosphorylation of serine or threonine to bind to 14-3-3 domains^65^. Three 14-3-3-binding motifs overlapped with regulated phosphorylation sites in the NELFE subunit of the negative elongation factor (NELF) complex (Figure 5B). We have previously shown that UV-C causes p38-dependent phosphorylation of the transcriptional regulator NELFE with consequent binding to 14-3-3 proteins^7^. We therefore went on to experimentally validate whether the NELFE - 14-3-3 interaction constitutes a general feature of the DDR. Indeed, endogenous NELFE co-immunoprecipitated much more with tagged 14-3-3 after HU, ETO and AsO_2_ treatment compared to treatment with DMSO, indicating that this interaction is required to regulate transcription upon a broader spectrum of stress responses (Figure 5C). Phosphorylation of histone H3.1 on serine 28, observed in the context of condensation of mitotic chromosomes and signal-induced transcription in response to mitogens or extracellular stressors, has been shown to trigger binding to 14-3-3 proteins^66,67^. We further show by co-immunoprecipitation that this interaction is lost specifically after ETO treatment (Figure 5C), in line with the strong dephosphorylation observed upon DSB induction (Figure 5B). Similarly, we establish that the interaction of transcriptional regulator LDB1 with 14-3-3 proteins decreased in AsO_2_-treated cells compared to other treatments (Figure 5C). A degradation motif, or degron, is a short sequence on a target protein that, upon recognition by a ubiquitin E3 ligase, leads to its ubiquitylation and degradation. We identified 11 degrons containing DNA damage-responsive phosphorylation sites, the majority of which are known substrates of the SCF (Skip1-Cullin-Fbox) ubiquitin ligase complex (Figure S5A). This subset of proteins included members of NF-kB pathway (NFKB2, NFKBIE), translation regulators (EEF2K, PCDC4, PRKRA) and the ubiquitin E3 ligase UHRF1. In addition, we found 12 trafficking SLiMs whose phosphorylation was modulated by DNA damage, including several nuclear localization sequences (HNRNPA1, NCPB1, MICAL3, SRF, TARDBP, XRCC1) (Figure S5B), indicating that nuclear localization of these proteins is dynamically regulated during the DDR.

**Figure 5.**
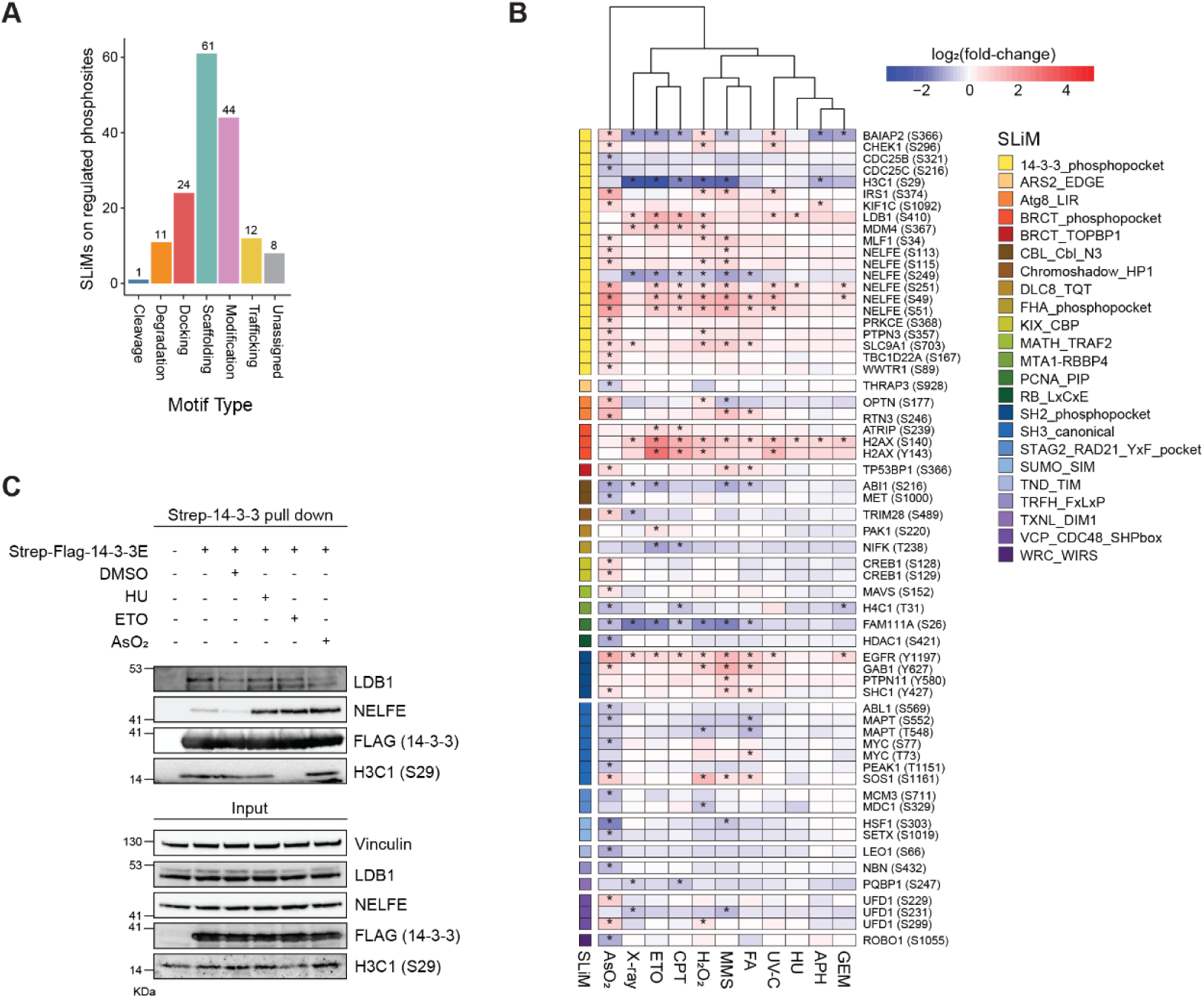
DNA damage-induced phosphorylation sites regulate SLiM-based interactions. A. Bar graph showing the number of SLiMs of each motif type harbouring significantly regulated (FDR ≤ 0.05; absolute fold-change > 1.5) phosphorylation sites. B. Heatmap showing significantly regulated phosphorylation sites overlapping with SLiMs of the Scaffolding type. Phosphosites are annotated with the respective SLiM name. C. Representative Western blot images after Strep-14-3-3 pulldown from whole-cell lysates of U2OS cells treated with DMSO, HU, ETO or AsO_2_ at the previously determined concentrations for 2h (n=1). Cells were transfected with a Strep-FLAG-14-3-3-E or an empty Strep-FLAG vector.

### Phosphorylation of proteasome-associated ubiquitin E3 ligase UBE3A by ATM/ATR is part of the core DNA damage response

To gain insight into how different DNA-damaging treatments affect proteome quality control pathways, we investigated the association of differentially phosphorylated proteins with specific branches of the proteostasis network (annotated by the Proteostasis Consortium)^68^. The highest proportion of differentially regulated proteins across all treatments belonged to the ubiquitin-proteasome system (UPS), in line with the well-described roles of the UPS and ubiquitylation in the DDR (Figure 6A). We found 106 ubiquitin E3 ligases and 32 deubiquitylating enzymes (DUBs) to be differentially phosphorylated upon DNA damage (222 and 68 phosphorylation sites, respectively): among these, enzymes with known roles in DNA repair pathways, such as BRCA1 (17 sites), TRIM28 (13 sites), TRIP12 (8 sites), USP36 (7 sites), and USP1 (4 sites) were most prominently regulated (Figure 6B, S6A and B). Notably, the core response to DNA damage and replication stress, highlighted by our clustering analysis (phosphorylation sites induced by all treatments, independently of AsO_2_ - cluster 5), included five sites on UPS factors - S368, S440 and S218 on ubiquitin E3 ligases RAD18, TRIM28 and UBE3A, respectively, S663 on DUB USP37, and S784 on the AAA-ATPase VCP (also known as p97). Thirteen additional phosphorylation sites were regulated in at least 8 out of 11 treatments, including phosphorylation sites on ubiquitin E3 ligases ARIH2, BRCA1, DTX3L and RNF181, the DUB USP28, and the proteasome subunit PSMD4 (Figure 6C). Notably, S218 on UBE3A is found within an S/T-Q motif, suggesting direct regulation of the proteasome by DNA damage signaling (Figure 6D). UBE3A phosphorylation on S218 after HU treatment and topoisomerase II inhibition was reported to be mediated by ATM/ATR in HEK293T cells^69^. In line with these findings, western blotting with an antibody against the phosphorylated S/T-Q motif confirmed the increased phosphorylation of GFP-tagged UBE3A after replication stress, which was dependent on ATM/ATR activity (Figure 6E). Overexpression of the UBE3A phosphorylation-dead mutant (GFP-UBE3A-S218A) was sufficient to abolish the signal of the phospho-S/T-Q antibody, substantiating that this residue is a *bona fide* target of the DDR kinases (Figure 6F). Considering the widespread phosphorylation of S218 on UBE3A upon different types of DNA damage, we investigated its functional relevance. We found that knockdown of UBE3A resulted in the decreased amount of phosphorylated H2AX after ETO treatment, suggesting a role of UBE3A in regulation of the DDR signaling cascade (Figure 6G). This decrease was rescued by the re-expression of the wild type UBE3A, but not by the phospho-dead UBE3A (Figure 6G, S6D). Notably, UBE3A knockdown cells or cells expressing the phospho-dead UBE3A mutant were more sensitive to HU and ETO in colony-forming assays (Figure 6H, I, S6D). In conclusion, UBE3A phosphorylation by ATM/ATR on a single site renders cells more resilient to genotoxic stress. UBE3A has been shown to ubiquitylate the proteasome receptor RPN10 (also known as PSMD4)^70^, as well as DNA repair factors^71^ and cell cycle regulators^72,73^. Therefore, UBE3A is likely to globally influence nuclear protein degradation in response to genotoxic stress through two complementary mechanisms: adjusting proteasome activity through interaction with RPN10 and facilitating the degradation of its specific substrates.

**Figure 6.**
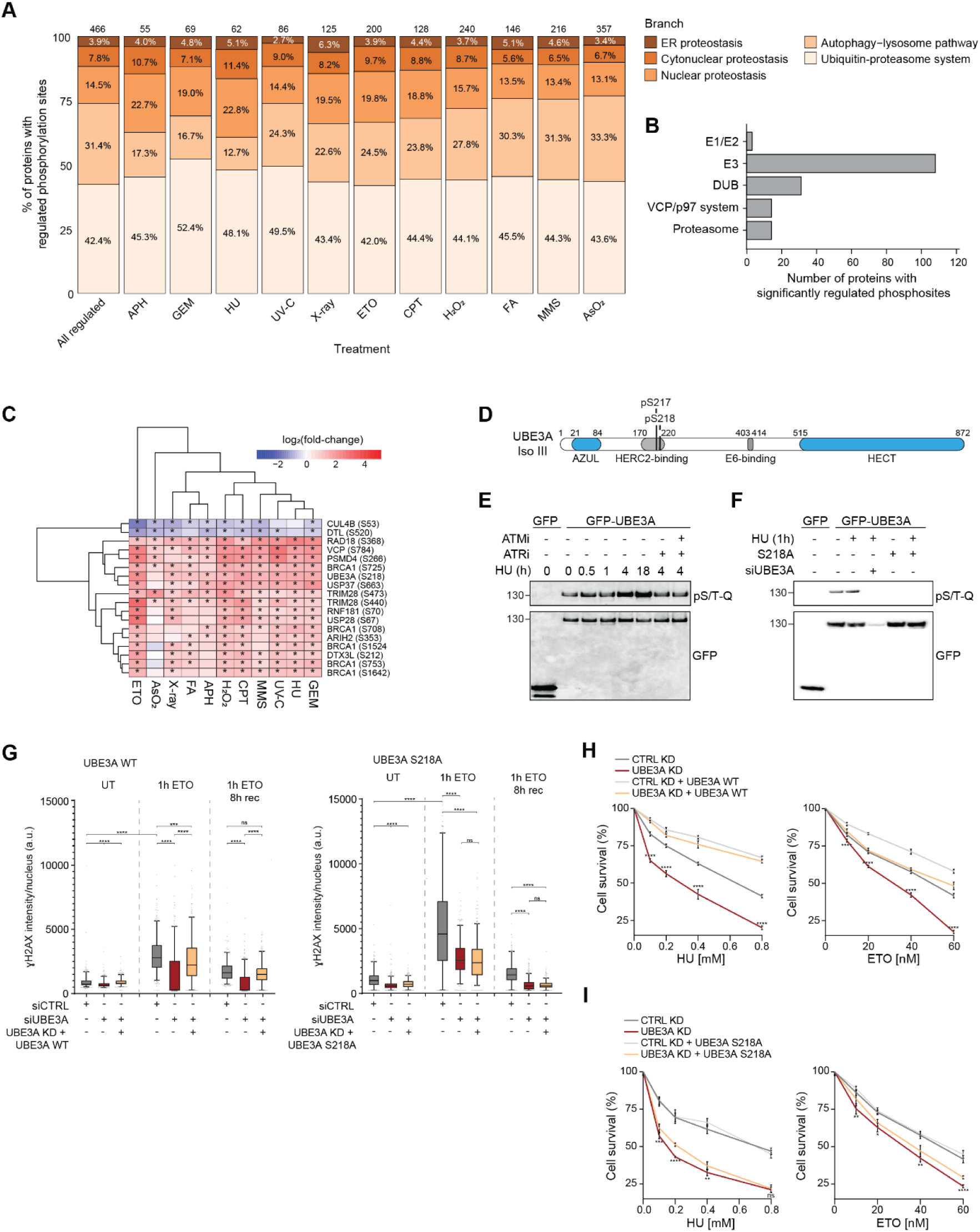
Phosphorylation of proteasome-associated ubiquitin E3 ligase UBE3A by ATM/ATR is a novel event in the core DNA damage response. A. Bar graph showing the percentage of differentially phosphorylated proteins (proteins harboring at least one significantly regulated phosphorylation site) associated with different branches of the proteostasis network, for each treatment. B. Bar graph showing the number of differentially phosphorylated proteins (proteins harboring at least one significantly regulated phosphorylation site) associated with different branches of the ubiquitin-proteasome system. C. Heatmap showing phosphorylation sites on members of the ubiquitin-proteasome system that are significantly regulated in response to at least 8 out 11 DNA damage treatments. D. Domain schematics of UBE3A. Phosphorylation sites S217 and S218 are indicated. E. Representative Western blot images after GFP-UBE3A pulldown from whole-cell lysates of doxycycline-inducible U2OS:GFP-UBE3A cells treated with doxycycline (1µM) and HU (2mM) for the indicated time-points, with or without ATM/ATR inhibitors (n=3). F. Representative Western blot images after GFP-UBE3A or GFP-UBE3A-S218A pull-down from whole-cell lysates of doxycycline-inducible U2OS:GFP-UBE3A or U2OS:GFP-UBE3A-S218A cell lines treated with doxycycline (1µM) and HU (2mM) for the indicated time-points. siRNA-mediated depletion of UBE3A was carried out for 48h prior to treatment with HU (n=3). G. Box plot displaying the levels of nuclear γH2AX in doxycycline-inducible U2OS:GFP-UBE3A (left) or U2OS:GFP-UBE3A-S218A (right) cell lines upon siRNA-mediated depletion of UBE3A and treatment with doxycycline (1µM) and ETO (10µM) for the indicated time-points. (*P-value < 0.05, **P-value < 0.01, ***P-value < 0.001, ****P-value < 0.0001) (n=3). H. Colony-forming assays using a doxycycline-treated U2OS:GFP-UBE3A cell line upon siRNA-mediated depletion of UBE3A and treatment with HU or ETO. Cells were treated at the indicated concentrations for 24 hours before, then allowed to recover for 10 days (n=3). I. Colony-forming assays using a doxycycline-treated U2OS:GFP-UBE3A-S218A cell line upon siRNA-mediated depletion of UBE3A and treatment with HU or ETO. Cells were treated at the indicated concentrations for 24 hours before, then allowed to recover for 10 days (n=3).

## Discussion

Humans experience DNA damage, replication stress, and oxidative stress originating from environmental sources, therapeutic interventions such as chemotherapy, and intrinsic metabolic processes. Protein phosphorylation mediated by ATM/ATR kinases orchestrates the cellular DDR by regulating DNA repair, cell cycle checkpoints and chromatin organization^1,2^. Early phosphoproteomics studies identified putative substrates of ATM and ATR after DNA damage and demonstrated the intricacy and breadth of DNA damage signaling in humans^6,74^. Over the past decade, phenotypic genetic screens and interaction-focused proteomic approaches have accelerated the discovery of novel DDR components. However, these strategies are limited in their ability to uncover the regulatory logic that governs the DDR. Comparative phosphoproteomics, as demonstrated here, addresses this gap by enabling the assignment of regulated phosphorylation sites and proteins to specific DNA repair pathways based on their phosphorylation dynamics in response to distinct genotoxic stressors and the predicted upstream kinases that mediate these modifications. The core DDR response comprised phosphorylation events common to both replication stress- and DSB-inducing agents, largely targeting S/T-Q motifs recognized by ATM and ATR kinases. Notably, S/T-Q phosphorylation was enriched among DNA repair factors regardless of conservation level. *Bona fide* DDR factors such as MDC1, TP53BP1, and RIF1 contained both poorly and highly conserved sites, suggesting that the ATM/ATR regulatory complexity within human DNA repair networks has increased. Importantly, more than 90% of DNA damage–induced S/T-Q phosphorylation sites remain functionally uncharacterized, underscoring that the core regulatory mechanisms of the DDR are still far from fully understood.

In addition to core DNA repair and cell-cycle checkpoint proteins, we detected S/T-Q phosphorylation on numerous proteins that were either previously unlinked or only loosely associated with DNA repair. Notably, we identified 175 S/T-Q sites across more than 100 known or putative RNA-binding proteins (RBPs), including highly conserved sites on splicing factors such as ACIN1, HNRNPF, SCAF1, SCAF11, ZC3H8, and ZC3H18. RBPs participate in several aspects of the DDR, including damage sensing, recruitment of repair factors, and post-transcriptional gene regulation^75–77^. Differential phosphorylation of RBPs may function as a molecular code that directs their recruitment to specific DNA lesions; for example, phosphorylation sites on THRAP3, BCLAF1, ALYREF, and CWC27 showed stronger induction by replication stress–inducing agents than by DSB-generating treatments. Mapping such regulatory phosphorylation events and their treatment-specific responses will facilitate the discovery of new mechanistic links between RNA-processing factors and the DDR.

Interestingly, we also observed low-conservation S/T-Q sites on proteins involved in transcriptional regulation, chromatin remodeling, and the ubiquitin–proteasome system. Protein degradation is a central component of the DDR, and recent work has shown that phosphorylation can regulate the activity and substrate specificity of the proteasomal subunit PSMD4 following DNA damage^78^. Consistent with this, we found that DNA damage induces phosphorylation of both core and regulatory proteasome components—including PSMA1, PSMB10, PSMD2, and PSMD4—as well as proteasome-associated E3 ubiquitin ligases UBE3A and RNF181, suggesting a broader ATM/ATR-dependent regulation of proteasomal function.

In addition to the canonical ATM/ATR phosphorylation signature, we found that MAPK and ZAKα signaling dominates the cellular response to FA, MMS, and oxidative stress–inducing treatments. This indicates activation of broader cellular stress programs, including the ribotoxic stress response (RSR) and the integrated stress response (ISR), both of which depend on these signaling pathways and are typically triggered by RNA damage^79^. Together, these observations underscore the multifaceted nature of the cellular response to genotoxic stress and highlight extensive crosstalk among stress-response pathways.

Assigning functions to individual phosphorylation sites remains challenging for several reasons, including the technical difficulty of expressing phospho-dead or phospho-mimetic mutants at physiological levels. Notably, we found that more than one-third of DDR-induced phosphorylation events occur within phospho-clusters. This is particularly relevant in the context of stress-induced biomolecular condensates, where multi-site phosphorylation enhances multivalent protein–protein or intramolecular interactions^79,80^. Indeed, recent studies have demonstrated that key DNA repair factors such as MDC1 and 53BP1 undergo phase separation into biomolecular condensates following various forms of DNA damage^81–84^. Phospho-clustering and the formation of charge blocks have been implicated in the partitioning of transcription factors into distinct condensates^85^, in nuclear speckle formation^86^, and in the organization of the chromosome periphery during mitosis^87^. Moreover, the phase behavior of numerous proteins—including INCENP^49^ and EIF4B^88^—has been shown to be regulated by clustered phosphorylation events. Given that multi-site phosphorylation is a defining feature of the DDR, mechanistic and functional characterization of regulated phospho-clusters will require systematic approaches that integrate computational modeling with experimental analyses. Our data further suggest that mutating a single phosphorylation site is often insufficient to elicit measurable phenotypic changes, underscoring the importance of studying clusters rather than individual residues.

Taken together, our work offers a resource of DNA damage–regulated phosphorylation sites, and their behavior in response to distinct genotoxic stressors, that can guide future investigations into the regulatory mechanisms governing cellular stress responses.

## Supplementary figures and legends

**Supplementary Figure 1.**
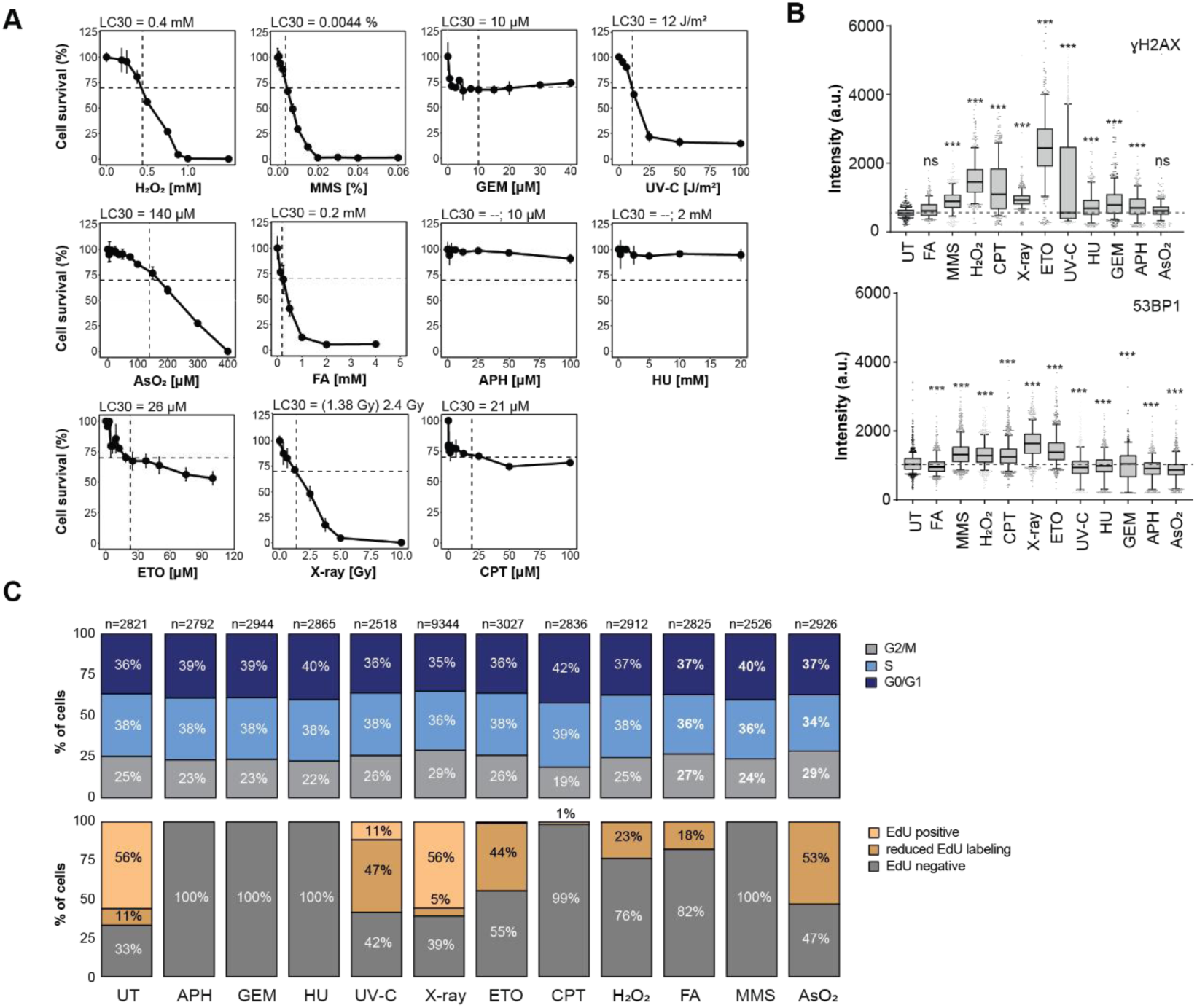
Defining the regulatory phosphoproteome of the DDR. A. Cell viability assay of U2OS cells 48 hours after treatment with increasing doses of DNA damaging agents for 2 hours. Cell viability was assessed using CellTiter-Blue viability assay or colony-forming assays (X-ray). Dose-response curves with the determined LC_30_ values are displayed (n = 3-6). Error bars indicate standard deviations. The LC30 dose could not be determined for APH and HU treatments. Instead, doses corresponding to complete inhibition of nucleotide reductase (2 mM HU) and DNA polymerase (10 µM APH), respectively, were chosen^89^ B. Box plots showing the quantification of DNA damage markers γH2AX or 53BP1 imaged with high-content fluorescence microscopy after 2-hour treatment of U2OS cells with the indicated genotoxic agent at the determined LC30. UV-C and X-ray-treated cells were allowed to recover for 2 hours post treatment. Significant pairwise comparisons were evaluated using one-way ANOVA followed by Dunnett’s test (*P-value < 0.01, **P-value < 0.001, ***P-value < 0.0001, compared to untreated -UT). C. Bar graph showing the cell cycle analysis of U2OS cells combining Hoechst staining and EdU incorporation after treatment with the determined LC_30_ for 2 hours or after 2 hour recovery time (UV-C and X-ray treatments). Images were acquired with a high-content fluorescence microscope. The percentages of cells in different cell cycle phases (upper panel) or with different levels of EdU incorporation (lower panel) are shown. The number of cells analyzed per condition is reported above each bar.

**Supplementary Figure 2.**
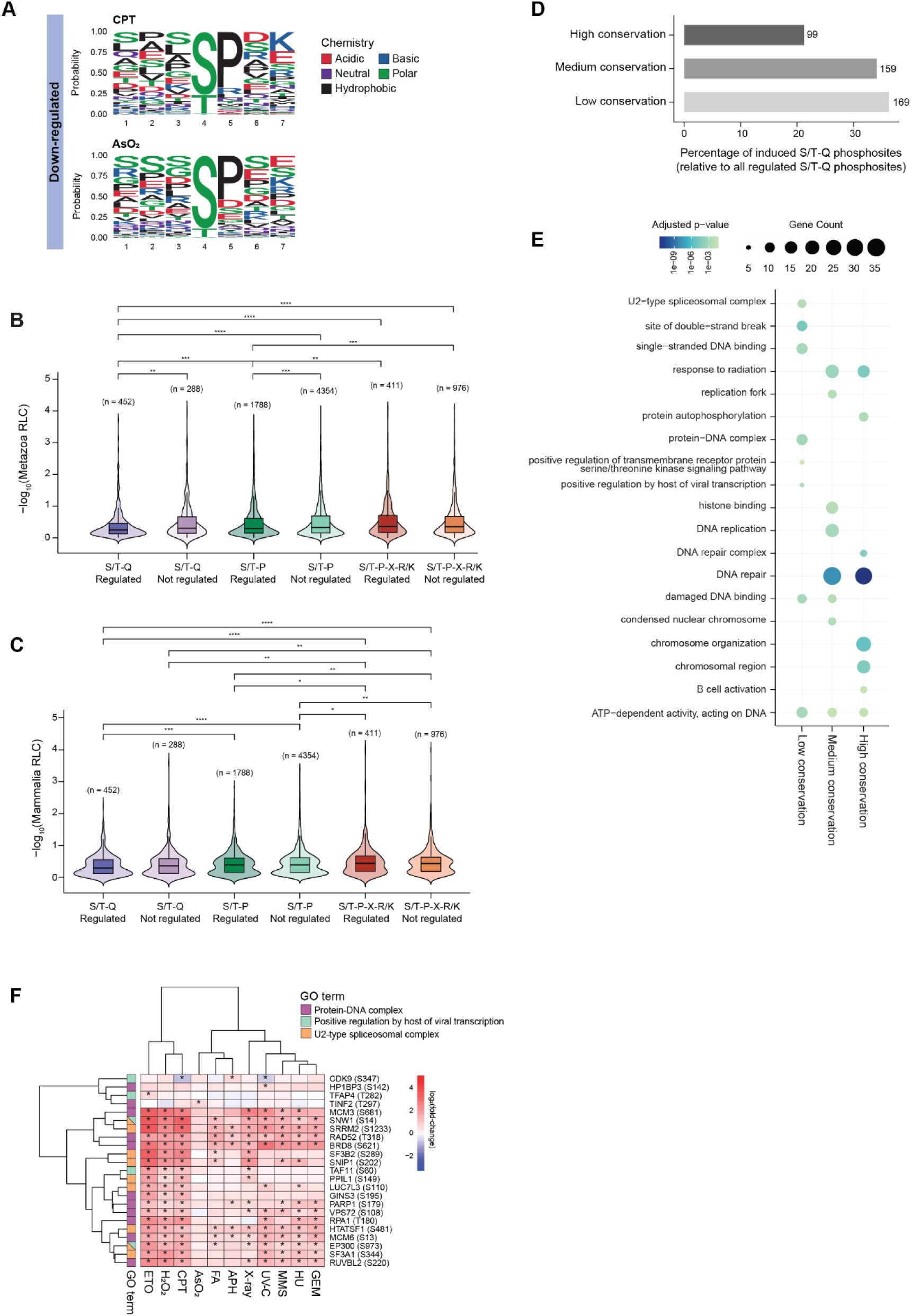
Protein kinase-substrate relationships in the DDR. A. Sequence motif analysis of treatment-specific phosphorylation sites gained (FDR ≤ 0.05; fold-change ≥ 1.5; upper panel) and lost (FDR ≤ 0.05; fold-change ≤ -1.5; lower panel) with iceLogo. Sequence windows of significantly changing sites were compared to all quantified phosphorylation sites. Selected treatments are shown. Amino acid chemistry is indicated (right). B. Violin plot displaying the -log10 transformed relative local conservation (RLC) score for metazoa for regulated (FDR ≤ 0.05; absolute fold-change > 1.5) or not regulated phosphorylation sites found in S/T-Q, S/T-P or S/T-P-X-R/K motifs. Statistical significance was assessed using Wilcoxon test with Benjamini-Hochberg correction (*P-value < 0.05, **P-value < 0.01, ***P-value < 0.001, ****P-value < 0.0001). C. Violin plot displaying the -log10 transformed relative local conservation (RLC) score for Mammalia for significantly regulated (FDR ≤ 0.05; absolute fold-change > 1.5) or not regulated phosphorylation sites found in S/T-Q, S/T-P or S/T-P-X-R/K motifs. Statistical significance was assessed using Wilcoxon test with Benjamini-Hochberg correction (*P-value < 0.05, **P-value < 0.01, ***P-value < 0.001, ****P-value < 0.0001). D. Bar graph showing the percentage of S/T-Q phosphorylation sites induced by DNA damage (FDR ≤ 0.05; fold-change ≥ 1.5) stratified by relative local conservation (RLC) score for metazoa (low conservation >= 0.65; medium conservation < 0.65 and > 0.35; high conservation =< 0.35). Absolute numbers of phosphorylation sites in each category are reported. E. Dot plot showing the comparative Gene Ontology term enrichment analysis for S/T-Q phosphorylation sites that are induced by DNA damage (FDR ≤ 0.05; fold-change ≥ 1.5), stratified by relative local conservation (RLC) score for metazoa. Only significant gene ontology terms (BH-adjusted p-value < 0.05) are shown. The first eight terms by fold enrichment per cluster are shown. Dot size corresponds to gene count. F. Heatmap of low conservation S/T-Q phosphorylation sites associated with selected GO terms and significantly changing (FDR ≤ 0.05; absolute fold-change > 1.5) in at least one condition.

**Supplementary Figure 3.**
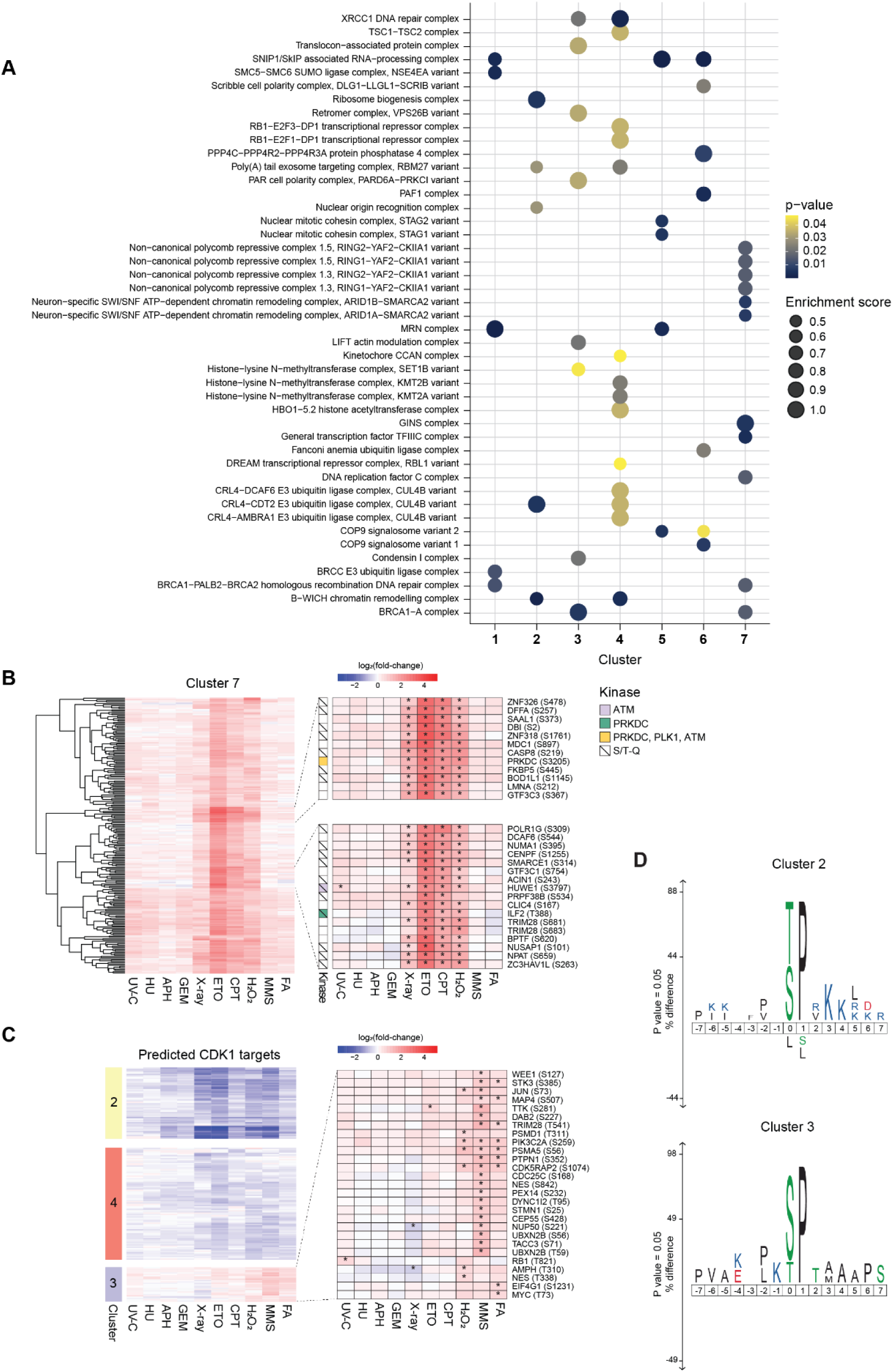
Specific, common, and pleiotropic responses of the phosphoproteome to DNA damage-inducing agents. A. Dot plot displaying the enrichment of protein complexes reported in the Complex Portal (EMBL-EBI) for each cluster. Only significantly enriched complexes (binomial test p < 0.05) with at least 2 differentially phosphorylated subunits are shown. B. Heatmap showing all cluster 7 phosphorylation sites (left), or zoomed-in panels with phosphorylation sites of interest (right). Phosphorylation sites are cross-referenced with the results of our KSEA analysis (CDKs, MAPKs, and PIKKs substrates) and S/T-Q sites are annotated. C. Heatmap showing CDK1 substrates (cross-referenced from our KSEA analysis) in clusters 1, 2 or 3 (left). A zoomed-in panel highlighting cluster 3 phosphorylation sites is presented on the right. D. Sequence motif analysis of treatment-specific phosphorylation sites gained (FDR ≤ 0.05; fold-change ≥ 1.5; upper panel) and lost (FDR ≤ 0.05; fold-change ≤ -1.5; lower panel) with iceLogo. Sequence windows of significantly changing phosphorylation sites were compared to all quantified phosphorylation sites. Amino acid chemistry is indicated (right).

**Supplementary Figure 4.**
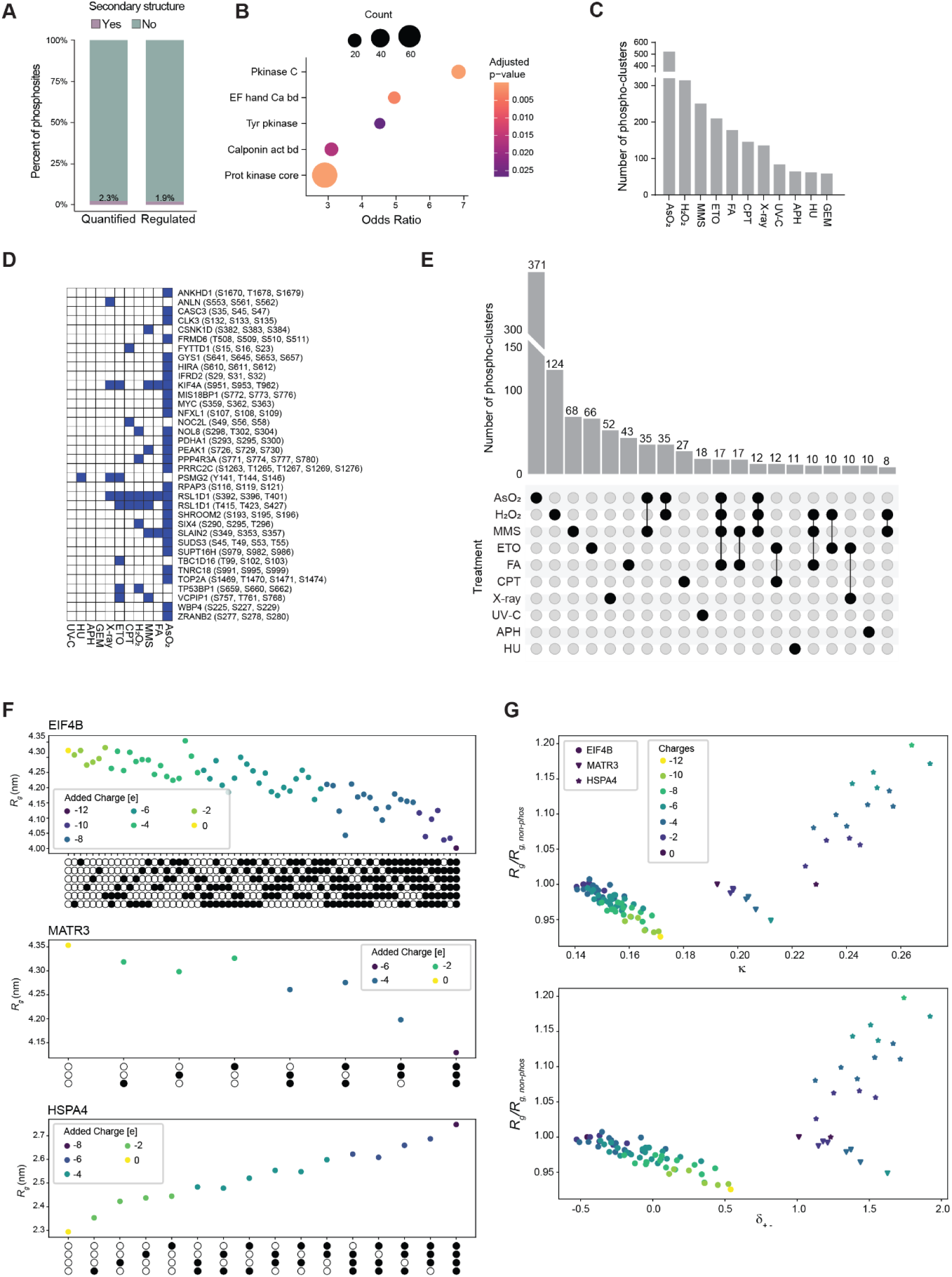
DDR-induced multi-site cluster phosphorylation regulates conformational dynamics of IDPs. A. Bar graph showing the percentage of either all quantified or significantly regulated (FDR ≤ 0.05; absolute fold-change > 1.5) phosphorylation sites found in disordered or structured regions. B. Dot plot showing the enrichment of protein domains harboring significantly regulated (FDR ≤ 0.05; absolute fold-change > 1.5) phosphorylation sites. C. Bar graph showing the number of phospho-clusters per treatment. Phospho-clusters were defined, separately for each treatment, as groups of three or more phosphorylated residues (independently of their regulation status) within a 10-amino acid stretch containing at least one treatment-responsive site. D. Heatmap showing the enrichment of phospho-clusters fully lost upon DNA damage treatments. E. Upset plot showing the overlap between phospho-clusters induced by different DNA damaging agents. The top 20 overlaps are shown. F. Scatter plots showing the mean radius of gyration plotted against the phosphorylation state of the protein. Each circle represents a phosphorylation site of the phospho-cluster and its phosphorylation state (empty: non-phosphorylated; filled: phosphorylated). G. Scatter plots showing the parameters κ (upper panel) or δ_+-_ (lower panel) plotted against the average radius of gyration, normalized by that of the corresponding unphosphorylated fragment, (R_g_/R_g,non-phos_,), with increasing charge added through phosphorylation.

**Supplementary Figure 5.**
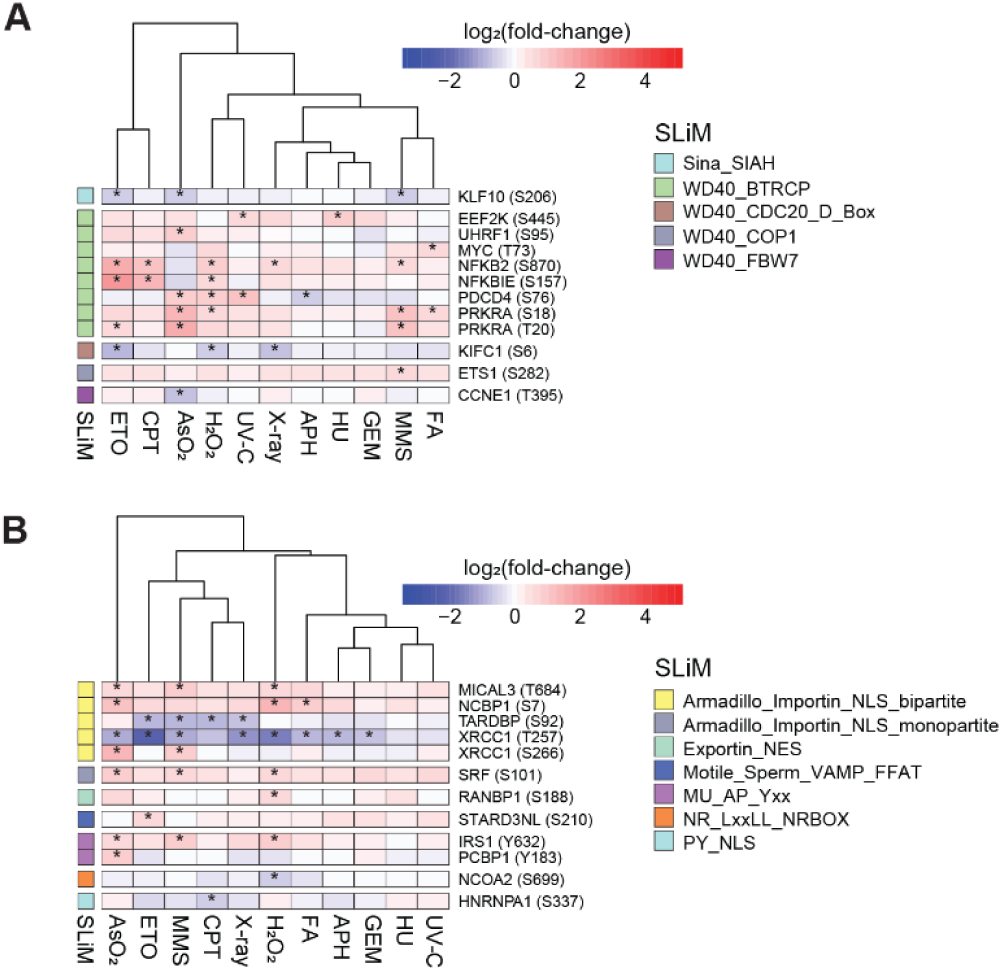
DNA damage-induced phosphorylation sites regulate SLiM-based interactions. A. Heatmap showing significantly regulated phosphorylation sites overlapping with SLiMs of the Degradation type. Phosphorylation sites are annotated with the respective SLiM name B. Heatmap showing significantly regulated phosphorylation sites overlapping with SLiMs of the Trafficking type. Phosphorylation sites are annotated with the respective SLiM name.

**Supplementary Figure 6.**
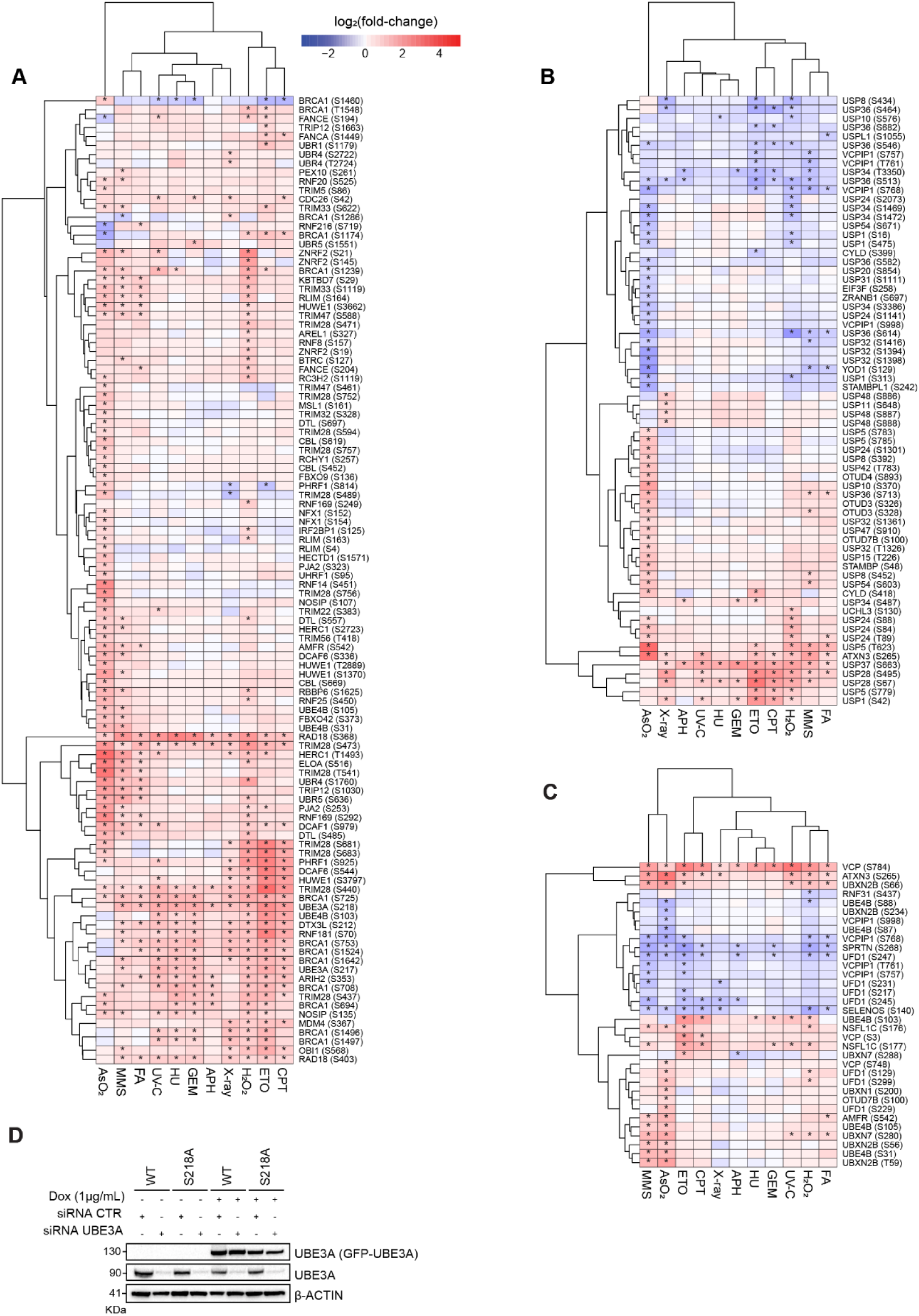
Phosphorylation of proteasome-associated ubiquitin E3 ligase UBE3A by ATM/ATR is a novel event in the core DNA damage response. A. Heatmap showing phosphorylation sites on E3 ubiquitin ligases induced (FDR ≤ 0.05; fold-change ≥ 1.5) by at least one treatment. B. Heatmap showing phosphorylation sites on deubiquitinating enzymes (DUBs) significantly changing (FDR ≤ 0.05; absolute fold-change > 1.5) in at least one treatment. C. Heatmap showing phosphorylation sites on VCP and VCP-associated proteins significantly changing (FDR ≤ 0.05; absolute fold-change > 1.5) in at least one treatment. D. Representative Western blot images after doxycycline-induced overexpression of siRNA-resistant UBE3A, wild type or phospho-dead (S218A) in U2OS cells.

## Supplementary dataset legends

**Supplementary dataset 1**: DNA damage phosphoproteome in response to 11 treatments in U2OS cells.

**Supplementary dataset 2**: Results of PepTools analysis.

**Supplementary dataset 3**: Significantly regulated induced S/T-Q phosphosites stratified by relative local conservation in metazoa and associated GO enrichment analysis.

**Supplementary dataset 4**: Results of SOM and k-means clustering with associated GO enrichment and complexome analysis.

**Supplementary dataset 5**: Results of domain enrichment analysis for phosphosites localized in structured regions.

**Supplementary dataset 6**: Results of phospho-clustering analysis.

**Supplementary dataset 7**: Phospho-clustering analysis after filtering and with total net charge calculation.

## Methods

### Cell cultivation

Human osteosarcoma cells (U2OS) and HEK293T were obtained from ATCC and cultured in D-MEM medium supplemented with 10% fetal bovine serum (FBS), 2 mM L-glutamine, and 100 U/ml Penicillin/Streptomycin (Thermo Fisher Scientific). Additionally, puromycin was added to stable cell lines to a concentration of 1 µg/µl (Merck). For SILAC experiments, cells were cultured in medium containing either L-arginine and L-lysine, L-arginine [^13^C6] and L-Lysine [^2^H4], or L-arginine [^13^C6,^15^N4] and L-lysine [^13^C6,^15^N2] (Cambridge Isotope Laboratories). All cells were grown at 37 °C in a humidified incubator at 5% CO_2_.

### Cloning

Expression vectors were produced by Gateway Cloning according to manufacturer’s instructions (Gateway LR Clonase II Enzyme mix; Thermo Fisher Scientific). Site-directed mutagenesis was used to remove mutations and to obtain an siRNA-resistent copy of UBE3A by introducing nucleotide wobbles at the siRNA target site^90^ Mutation of serine to alanine at position 218 was introduced in the same fashion. Lentivirus destination vector plix-GFP(c-term) was obtained through the introduction of a GFP-tag from pcDNA-DEST47 (Thermo Fisher Scientific) into the plix-402 (Addgene) background by CPEC cloning.

### Transfection of cells

Overexpression of vectors was performed using polyethylenimine (PEI, Polyethylenimine, 24885-2) with a DNA:PEI ratio of 1:3. Knockdown by siRNA transfection of cells was carried out using Lipofectamine RNAiMAX (Thermo Fisher Scientific) according to the manufacturer’s instructions.

### Stable cell line production by lentivirus transduction

For lentivirus production, HEK293T cells were co-transfected with an expression plasmid (pliX-UBE3A-GFP), along with the packaging plasmids psPAX2 and pMD2.G (Addgene) in a 4:3:2 w/w/w ratio. The supernatant was collected 72 hours post transfection, filtered through a 0.45 µm filter, mixed with the same volume of fresh medium, and supplemented with polybrene (Sigma-Aldrich) to a concentration of 8 µg/µl. Subsequently, cells were transduced by incubation with the conditioned supernatant. Stable cells were selected 72h post transduction by the addition of 2 µg/µl puromycin for 5 days.

### Colony-forming assay

Cells were transfected with the respective siRNAs. In the case of doxycyclin-inducible cell lines, doxycycline was added to a concentration of 1 µg/µl. The next day, 4,000 to 12,000 cells were re-seeded into 6-well plates. 72 hours post transfection, the cells were irradiated or treated with DNA damage-inducing agents for a period of 24 hours, followed by two PBS washes and medium exchange. Surviving cell colonies were stained with crystal violet solution 10-14 days post treatment and counted under the microscope. Three biological replicates with three technical replicates each were measured per condition. LC_30_ values were calculated by regression analysis in GraphPad prism (Dotmatics). LC_30_ values equivalent to the ones determined by CellTiter-Blue were determined by calculating scaling factors after carrying out both assays upon UV-C treatment.

### Cell viability assay

2,500 cells per well were seeded onto 96-well plates in a volume of 100 µl. Cells were treated with DNA damage-inducing agents for 2 hour, or irradiated, the next day, followed by medium exhange. Cell viability was tested 72 hours post treatment by adding 20 µl of CellTiter-Blue Cell Viability solution (Promega) for 3 hours. Bioluminescence was measured on a Tecan Spark Plate Reader with the following settings: excitation wavelength 560 nm, emission wavelength 590 nm, number of flashes 25, and integration time 20 μs. LC_30_ values were calculated by regression analysis in GraphPad prism.

### Immunofluorescence and confocal microscopy

For microscopy experiments, 20,000 cells were seeded into 96-well cell carrier plates. The next day, cells were treated according to the experimental setup. After two PBS washes, cells were fixed with 4% PFA solution for 20 min at RT, washed three times with PBS, and incubated in blocking buffer for 1 hour. Primary antibodies were diluted according to the manufacturer’s instructions in blocking buffer and incubated with the cells overnight at 4 °C. Cells were washed 3 times with PBS-T, followed by incubation with secondary antibody diluted in blocking buffer and in combination with Hoechst 33342 (2 µg/µl; Thermo Fisher) for 1 hour at RT. Cells were washed three times with PBS-T and stored in PBS until the measurement. Images were acquired on the Opera Phenix High Content Screening System using a 40X water objective lens. Around 30 fields per well with 5-10 z-planes separated by 0.5 μm were obtained. Image analysis was performed in Harmony High-Content Imaging and Analysis Software. Significance values were calculated in GraphPad (one-way ANOVA followed by Dunnett’s test).

### Cell-cycle profiling

Cells were grown to 80% confluence, and EdU was added to a concentration of 10 µM for 1 hour. For the analysis by flow cytometry, the cells were harvested and washed with PBS and fixed for 10 min in 4% PFA. Click reaction was carried out by resuspension of cells in 3480 µl H_2_O, and sequential addition of 400 µl sodium L-ascorbate (100 mM), 40 µl Alexa Fluor 647 Azide (10 mM), and 80 µl copper sulfate (100 mM). The clique reaction was incubated for 30 min at RT. Afterward, DNA was stained with either Hoechst 33342 (1 µg/ml) or 7-Aminoactinomycin D (7-AAD, 1:100; Thermo Fisher). Cells were analyzed on the BD LSRFortessa SORP cell analyzer (7-AAD: BL488nm, 610/20 nm; Alexa Fluor 647: RL635nm, 660/20; Hoechst 3342: 355nm, 450/50). For cell cycle analysis by microscopy, click reaction, Hoechst staining and subsequent measurement on the Opera Phenix High Content Screening System (PerkinElmer) were carried out as described above.

### Cell lysis

Cells were washed twice with ice-cold PBS, placed on ice, and directly lysed in the plates by addition of modified RIPA buffer (50 mM Tris-HCl pH 7.5, 150 mM NaCl, 1 mM EDTA, 1% NP-40, 0.1% Sodium-deoxycholate) supplemented with protease (Complete protease inhibitor cocktail tablets, Roche Diagnostics) and phosphatase inhibitors (1 mM sodium orthovanadate, 5 mM β-glycerophosphate, 5 mM sodium fluoride, Sigma-Aldrich). For samples prepared for Western blotting, NaCl was added to a concentration of 600 mM and sonicated. The lysates were cleared by centrifugation at 16.000 x g for 15 min at 4 °C, and protein concentrations were measured using the QuickStart Bradford Protein assay (Biorad).

### SDS-PAGE and Western blotting

LDS buffer containing protein samples were separated on NuPAGE Bis-Tris gels with a 4-12% gradient (Thermo Fisher Scientific) at 150 V in the MOPS SDS running buffer. Proteins were transferred onto a 0.45 µm nitrocellulose membrane (Sigma Aldrich) in a transfer buffer using the Xcell II Blot-Modul (Thermo Fisher Scientific) with 30 V for 115 min. All subsequent steps were carried out on a shaking platform and between the steps, the membrane was washed 3 times with a PBS-T buffer (1x PBS, 0.1% Tween-20). First, the protein transfer was checked by Ponceau S (Thermo Fisher Scientific) staining of the membrane. Next, the membrane was incubated in blocking buffer (5% FBS in PBS-T) for 1 hour at RT, followed by incubation with primary antibody diluted in blocking buffer overnight at 4 °C. Lastly, the membrane was incubated with a secondary antibody coupled to horseradish peroxidase (1:5000 in blocking buffer) for 1 hour at RT. Signal detection was carried out in the ChemiDoc imaging system (Bio-Rad) in combination with the SuperSignal West Pico Chemiluminescent Substrate (Thermo Fisher Scientific).

### Pull-down using GFP-Trap agarose

Cell lysates were prepared as described above. 20 µl of pre-equilibrated GFP-trap (ChromoTek) beads were added to 1 mg of lysate and incubated for 1 hour at 4 °C with rotation. The beads were washed 3 times with a modified RIPA buffer supplemented with protease and phosphatase inhibitors, and an excessive buffer was drained with a syringe. The dried beads were resuspended in 35 µl of 2x NuPAGE LDS sample buffer (Thermo Fisher Scientific) supplemented with 1 mM dithiothreitol (DTT) and incubated at 70 °C for 10 min before the supernatant was applied for SDS-PAGE. For bait protein isolation, GFP-PDs were washed 3 times with 8M urea in PBS and twice in with RIPA buffer, before the proteins were eluted in LDS buffer.

### In-solution protein digest

Cell lysates were prepared as described above. Proteins were precipitated in fourfold excess of ice-cold acetone and incubated overnight at -20 °C, followed by pelleting by centrifugation for 5 min at 1000 x g. Subsequently, the pellets were re-dissolved in denaturation buffer to a concentration of 2-8 µg/µl. Cysteines were reduced with 2 mM DTT in the dark and alkylated with 10 mM CAA for 40 min respectively. For protein digestion, sequencing grade-modified trypsin (Sigma-Aldrich) was added in a 1:75 ratio overnight. Protease digestion was stopped by the addition of TFA to a concentration of 0.5% and the samples were incubated for 30 min at 4 °C. Formed precipitates were removed by centrifugation for 10 min at 4000 x g. To purify and concentrate the peptides, reversed-phase Sep-Pak C_18_ cartridges (Waters) were prepared by washing them once with ACN and three times with 0.1% TFA. After peptide samples were loaded, the columns were washed three times with water, dried, and stored at 4 °C.

### TMT labeling

Peptides were eluted from SepPak columns in 50% ACN, fully dried in a vacuum concentrator at RT, and resuspended in labeling buffer (100 mM HEPES, pH8.5). Concentrations were determined with the Nanodrop (Thermo Fisher) at 280 nm and adjusted to 5 µg/µl. For each labeling reaction, 100 µg of peptides were mixed with an equal amount of TMT label (20 µg/µl, TMT-10plex + TMT11-131C, Thermo Fisher) and incubated for 1 hour at 25 °C. Labeling reactions were quenched by the addition of hydroxylamine to a final concentration of 0.4% and incubated for 20 min at 25 °C. Subsequently, the peptide samples were diluted in 0.1% TFA, reducing the ACN concentration below 3%. At this point, peptide labeling was assessed as previously described (Zecha et al. 2019). Fully labelled TMT samples were pooled and purified on Sep-PAK columns as described above.

### Phosphopeptide enrichment

TMT labeled peptides were eluted from Sep-PAK columns in 2 ml 50% ACN and acidified to a concentration 6% TFA. 1 mg of peptides were mixed with 2 mg of TiO_2_ beads (10 μm, GL Sciences Inc) and incubated for 1 hour at RT with rotation. Beads were washed twice with 1 ml binding buffer (50% ACN, 6% TFA) and twice 1ml wash buffer (50% ACN, 0.1% TFA). They were loaded onto a StageTip made with 1 layer of a C_8_ 47 mm extraction disk (Empore) and dried by centrifugation at 500 x g. Phosphorylated peptides were eluted with 100 µl of elution buffer 1 (5% NH_4_OH), followed by 100 µl of elution buffer 2 (10% NH_4_OH, 25% ACN) by centrifugation at 400 x g. The eluted peptides were vacuum concentrated at 45 °C for 20 min to remove NH_4_OH. pH was adjusted to < pH 2 with TFA and peptides were fractionated by Micro-SCX.

### Fractionation by SCX and desalting of peptide samples

Peptide fractionation and purification was carried out using stop-and-go-extraction tips (StageTips)^91^. For fractionation, peptide samples were loaded onto SCX StageTips and sequentially eluted by increasing pH (pH 2.5-11, 40 mM acetic acid, 40 mM boric acid, 40 mM phosphoric acid). Eight and twelve fractions were collected for peptide and phosphopeptide samples respectively. Fractionated samples were vacuum centrifuged to remove ACN, acidified, and desalted on C_18_ StageTips. Samples were eluted in 50% ACN, 0.1% TFA and vacuum concentrated at 45 °C to completion. Finally, samples were reconstituted in 0.1% FA, 2% ACN.

### MS analysis

Peptide fractions were analyzed on quadrupole Orbitrap mass spectrometers (Exploris 480) equipped with a UHPLC system (EASY-nLC 1200, Thermo Scientific). Peptide samples were loaded onto C_18_ reversed-phase columns (length 55 cm, inner diameter 75 μm; Reprosil Pur 1.9 μm, Dr Maisch) and eluted in a linear gradient from 8 to 40% acetonitrile containing 0.1% formic acid in 105 min. The mass spectrometer was operated in DDA mode, automatically switching between MS1 (60k resolution, 300% AGC target) and MS2 acquisition (15k acquisition, 100% AGC target). Survey full-scan MS spectra (m/z 350–1650) and fragment spectra were acquired in the Orbitrap^92–94^. MS1 and MS2 maximum injection times were set to 28 ms respectively. The isolation window was set to 0.8 m/z. The 15 most intense ions were sequentially isolated and fragmented by HCD (25 normalized collision energy). TurboTMT was switched on, while advanced peak determination was deactivated^95,96^. Peptides with unassigned charge states, as well as with charge states less than +2 were excluded from fragmentation.

### MS peptide identification

Raw data files were analyzed using MaxQuant (version 2.6.5.0)^97^. Parent ion and MS2 spectra were searched against a database containing 96,788 human protein sequences obtained from the UniProtKB released in March 2025 using the Andromeda search engine^98^ Spectra were searched with a mass tolerance of 6 ppm in MS mode, 20 p.p.m. in HCD MS2 mode, strict trypsin specificity, and allowing up to three miscleavages. Cysteine carbamidomethylation was searched as a fixed modification, whereas protein N-terminal acetylation, methionine oxidation, and phosphorylation of serine, threonine, and tyrosine were searched as variable modifications. Spectra were filtered for precursor ion fractions (PIF) > 0.75. The dataset was filtered based on posterior error probability to arrive at a false discovery rate below 1% estimated using a target-decoy approach^99^

### Statistical analysis of MS data

Proteins/Peptides identified from the reversed database, identified by site, or identified as potential contaminants were excluded, as were proteins with no unique or less than two identified peptides. The cut-off criterion for the localization probability of each phosphosite was ≥ 0.75 (= class I sites). For the phosphorylation site analysis, the lowest available underscore intensity entries were used. Furthermore, protein and peptide intensities were normalized to correct for loading mistakes and batch effects. Isotope corrected intensities were first divided by the corresponding reference intensity separately for each batch (reference normalization). In a second step, intensities in each sample were divided by the sample mean/median (mean/median centering). Statistical analysis and data visualization was performed using the R software environment (v4.2.3). Moderated *t*-tests corrected for multiple hypothesis testing (Benjamini-Hochberg) were calculated using the limma package^100^, including only peptides/proteins measured in at least two replicates.

### Phospho-cluster identification

Phospho-cluster identification was performed separately for each treatment on proteins harboring significantly changing phospho-sites. Phospho-sites located within a 10 amino-acid window along the protein sequence were grouped using hierarchical clustering and significance was assessed by comparing to a null distribution generated from 1,000 iterations in which the same number of phospho-sites was randomly sampled from all phosphorylatable residues (S/T/Y) in the protein. Phospho-clusters with FDR =< 0.01, composed of two phospho-sites, or only of phospho-sites undergoing non-significant changes in phosphorylation status were filtered out. Phospho-cluster charge was calculated by summing up the charge of single charged residues (K, R = positive charge; D, E = negative charge; phospho-site = double negative charge).

### Other bioinformatic analysis

Kinase prediction analysis was perfomed using the KSEA algorithm (KSEAapp) with updated known kinase-substrate relations (PhosphoSitePlus, 04/2021) and a prediction NetworKIN cutoff of 5^26,101^. Sequence motif analysis was performed with IceLogo^102,103^. Dimensionality reduction by self-organizing maps (SOM) was performed with the kohonen R package^104^ and followed by k-means clustering (k=7). Gene Ontology (GO) enrichment analysis was performed with ClusterProfiler^105^ using the full phosphoproteomics dataset as background universe and the Benjamini-Hochberg method for p-value correction. Protein domain enrichment was performed with EnrichR using the full phosphoproteomics dataset as background universe^106^

### AlphaFold models

The structures of MATR3, HSPA4 and EIF4B were taken from the AlphaFold Database, predicted with the AlphaFold Monomer v2. ^54^. Parts of the structure with AlphaFold with very high (plDDT>90) or confident (70<plDDT<90) confidence score were assumed to be folded, other parts of the structure were assumed to be disordered. The disordered sections containing the phosphorylation sites were extracted for further analysis: residues 391-611 for EIF4B, residues 582 - 781 for MATR3 and residues 504-573 for HSP4A.

### Coarse-grained replica exchange stochastic molecular dynamics simulations

The disordered sections containing phosphorylation-rich regions (EIF4B 391-611, MATR3 582-781, HSP4A 504-573) were simulated by coarse-grained (CG) replica exchange stochastic molecular dynamics simulations. The fragment was simulated by varying the phosphorylation extent of each phosphocluster. Accordingly, each phosphorylation site was systematically phosphorylated, considering all possible combinations (i.e. phosphorylating individual sites, or pairs, triads, tetrads, pentads or sextads of them), yielding a total of 8 (MATR3), 16 (HSP4A) and 64 (EIF4B) possible phosphorylation conditions (one of them being the fully dephosphorylated case). Simulations were carried out using CALVADOS 2^107,108^, as previosuly described^49^. Following this method, each amino acid of the fragment was modelled as a CG bead. Beads were connected by harmonic springs (elastic constant 8368 kJ/mol and equilibrium distance 0.38 nm) and beads separated by at least two bonds also interacted via non-bonded electrostatic and short-range interactions. Electrostatics between bead pairs were described by the Debye-Hückel electrostatic potential, assuming a dielectric constant of the solvent ɛ_r_=80, and an ionic strength of 0.1 M at 300 K (which corresponds roughly to a Debye-Hückel length κ_DH_ of 0.97 nm). Unphosphorylated and phosphorylated aminoacid charges were taken from Teseu et al. ^107,108^. Short range interactions were modeled by means of the Ashbaugh-Hatch potential^109^, setting the strength of the interaction to ɛ=0.8368 kJ/mol, adjusted to consider the hydrophobicity of each amino acid according to the λ parameters taken from the Calvados model^107,108^. For the phosphorylated residues, the lowest λ value, corresponding to the highly hydrophilic Glu residue, was considered. The size of each amino acid was considered via the σ parameter taken from Regy et al.^110^ for the standard amino acids, and from Perdikari et al.^111^ for their phosphorylated versions. Electrostatic interactions were truncated at a cutoff of 4 nm (i.e. ∼4.12ᐧκ_DH_) and short-range interactions at a cutoff of 2.4 nm (i.e. ∼2.4ᐧκ_DH_). The initial CG bead positions were assumed to be the c-alpha atomic positions in the all-atom structure of the studied fragment predicted by AlphaFold. Each fragment was placed in a simulation box of dimensions (167.96001 nm)^3^, sufficiently large to accommodate any of the fragments in a fully stretched conformation. Throughout all the simulations the volume was maintained constant (NVT ensemble conditions). Phosphorylation was introduced by modifying the mass, charge, λ and σ of the target amino acids to be phosphorylated. Stochastic molecular dynamics were used to describe the dynamics of the CG beads. The stochastic dynamics integrator, available in the GROMACS package^112,113^ (version 2019), was used for this purpose. Equations of motion were integrated at discrete time steps Δt= 5 fs. For each amino acid, the friction coefficient was computed as the ratio between its mass (taken from Tesei et al.^107,108^) and the relaxation time τ_t_=50 ps. User-defined tabulated potentials were used to compute both short-range and electrostatic interactions.

To enhance conformational sampling, temperature replica-exchange was employed. Accordingly, 36 simulation replicas ran in parallel at different temperatures, allowing the swapping of coordinates between adjacent replicas every 5 ns. A temperature interval from 151.685 to 617.62 K was chosen,ranging from low values - where the fragments were expected to be compact - to high values - where they are thought to be mostly extended. The chosen reference temperatures were 151.685, 157.894, 164.357, 171.084, 178.087, 185.376, 192.964, 200.863, 209.084, 217.642, 226.551, 235.824, 245.477, 255.524, 265.983, 276.871, 288.203, 300.000, 312.280, 325.062, 338.367, 352.217, 366.634, 381.641, 397.262, 413.522, 430.449, 448.068, 466.408, 485.499, 505.371, 526.057, 547.589, 570.003, 593.334, and 617.620 K. The temperatures are exponentially spaced following the formula aᐧexp(bᐧT), with a and b chosen such that they covered the desired range and that room temperature (300 K) was in the middle of the interval. The total simulation time was 2 microseconds per replica. The first microsecond was discarded from the analysis as it was accounted as equilibration. In the main text, results obtained at 300 K were considered.

Different structural properties of the simulated protein sections were extracted from the simulations using GROMACS gmx tools^113^. The radius of gyration R_g_, a measure of the size of a protein, was calculated using the gmx gyrate tool, and normalized by the mean radius of gyration of the unphosphorylated protein for a size-independent comparison. The probability of forming a contact was estimated from the fraction of the simulation in which any aminoacid was in contact with any other aminoacid (i.e. below a cutoff distance of 0.8nm). Inter-residue distances were extracted using the gmx pairdist command.

Kappa is a measure of the extent of charge segregation in a sequence as defined by Das & Pappu^50^, who also showed that measures of protein size decrease as kappa increases, due to the long-range electrostatic attractions. Kappa, as well as fraction of charged residues (FCR) and net charge per residue (NCPR) were calculated using localcider v.0.1.12^114^. Kappa is influenced by sequence length and composition, so an alternative method of quantifying the charge segregation is using z(delta+-), which is also known to be elevated when protein size decreases^51,52^. z(delta+-) was calculated using Nardini v.1.1.1^51^.

## List of antibodies used in this study

| Protein name | Product number | Origin | Dilution |
| --- | --- | --- | --- |
| 53BP1 | MAB3802 | Millipore | 1:400 (IF) |
| β-Actin | A2228 | Sigma | 1:5000 (WB) |
| FLAG | F1804 | Sigma | 1:1000 (WB) |
| GFP | sc-9996 | Santa Cruz Biotechnology | 1:2000 (WB) |
| γH2AX | A300-081A-M | Bethyl | 1:1000 (IF) |
| LDB1 | 64994 | Cell Signaling | 1:1000 (WB) |
| NELFE | HPA007187 | Sigma | 1:1000 (WB) |
| Phospho-Histone H3 (Ser28) | 9713 | Cell Signaling | 1:1000 (WB) |
| pS/pT-QG | 6966 | Cell Signaling | 1:1000 (WB) |
| UBE3A | 7526 | Cell Signaling | 1:1000 (WB) |
| Vinculin | V9264 | Sigma Aldrich | 1:10000 (WB) |
| Goat Anti-Mouse Immunoglobulins-HRP | P044701-2 | Agilent | 1:5000 (WB) |
| Goat Anti-Rabbit Immunoglobulins-HRP | P044801-2 | Agilent | 1:5000 (WB) |
| Goat Anti-Mouse IgG; Alexa Fluor 488 | A11001 | Life Technologies | 1:1000 (IF) |
| Goat Anti-Rabbit IgG; Alexa Fluor 568 | A11004 | Life Technologies | 1:1000 (IF) |
| Goat Anti-Mouse IgG; Alexa Fluor 488 | A11008 | Life Technologies | 1:1000 (IF) |
| Goat Anti-Rabbit IgG; Alexa Fluor 568 | A11011 | Life Technologies | 1:1000 (IF) |

## List of siRNAs used in this study

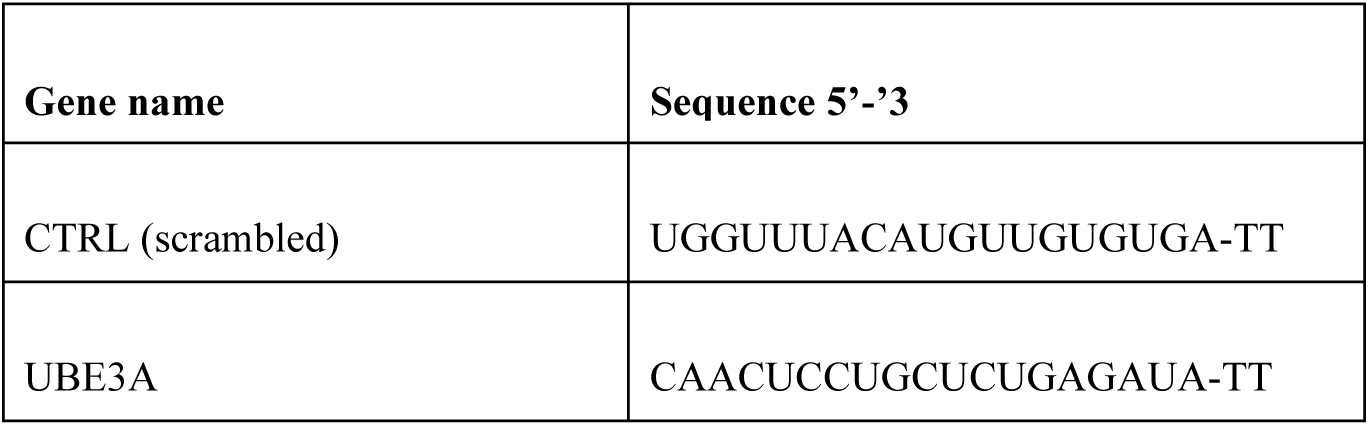

## List of oligos used in this study

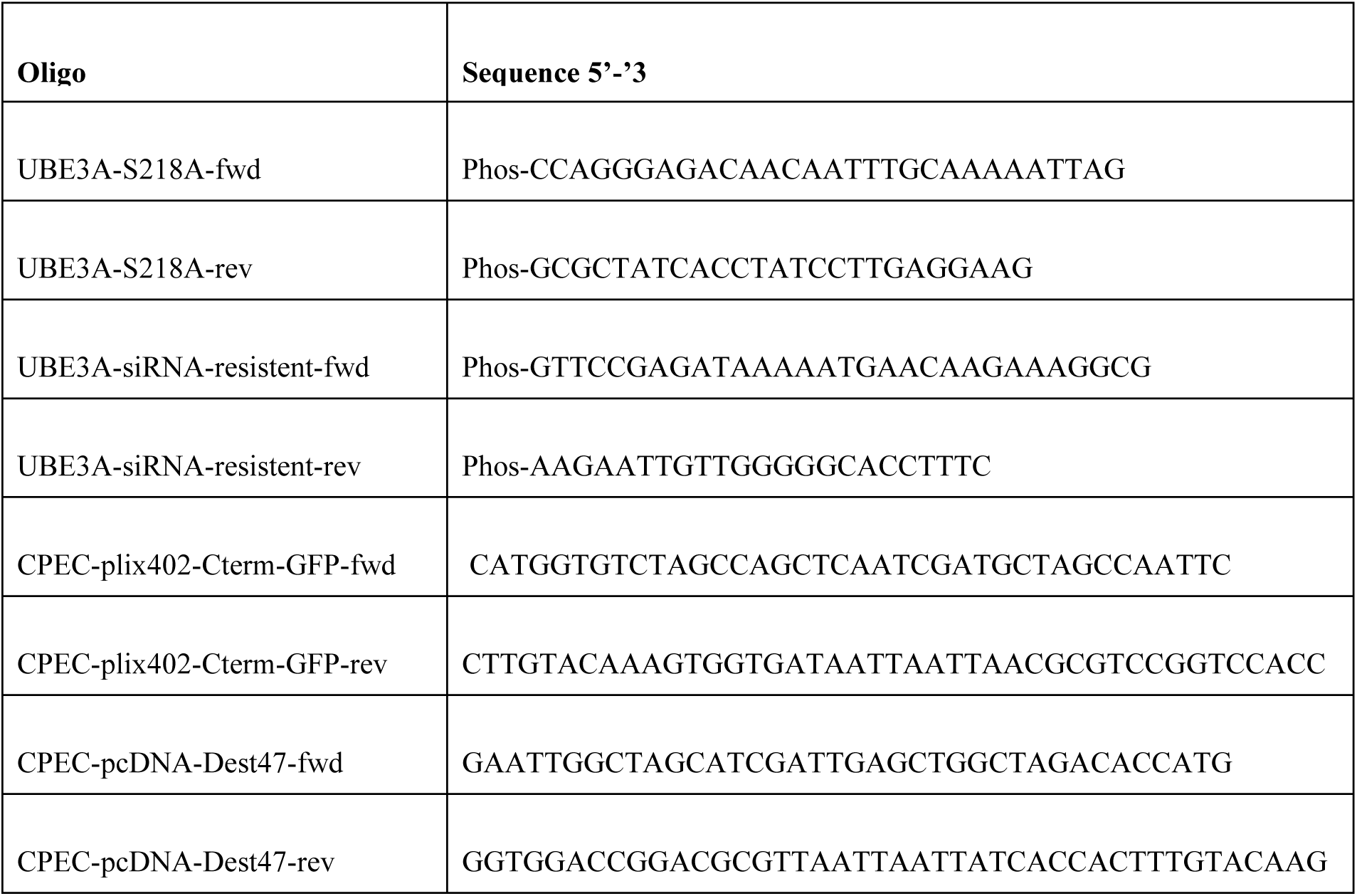

## List of plasmids used in this study

| Construct | Source |
| --- | --- |
| pCDNA-DEST47 | Life Technologies |
| pENTR221-UBE3A (isoform 3) | This study |
| pENTR221-UBE3A-S218A | This study |
| pMD2.G | Addgene |
| psPAX2 | Addgene |
| plix-402 | Addgene |
| plix-Cterm-GFP | This study |

## Acknowledgments

We thank Katharina Lepp for technical support, Amitkumar Fulzele and the IMB proteomics CF for help with mass spectrometry. Support from the IMB Microscopy, Flow Cytometry, Bioinformatics and Protein Production Core Facilities is gratefully acknowledged. The research for this project in the Beli lab is funded by the Deutsche Forschungsgemeinschaft (German Research Foundation, DFG) project-ID 464588647—SFB 1551 and project-ID 393547839—SFB 1361.

